# Single-Molecule Studies of Cognate and Near-Cognate Elongation in an *in vitro* Eukaryotic Translation System

**DOI:** 10.1101/2024.08.29.609187

**Authors:** Clark Fritsch, Arpan Bhattacharya, Martin Y. Ng, Hong Li, Philip C. Nelson, Barry S. Cooperman, Yale E. Goldman

## Abstract

The ribosome plays a central role in translation of the genetic code into amino acid sequences during synthesis of polypeptides. During each cycle of peptide elongation, the ribosome must discriminate between correct and incorrect aminoacyl-tRNAs according to the codon present in its A-site. Ribosomes rely on a complex sequence of proofreading mechanisms to minimize erroneous selection of incorrect aminoacyl-tRNAs that would lead to mistakes in translation. These mechanisms have been studied extensively in prokaryotic organisms, but eukaryotic elongation is less well understood. Here, we use single-molecule fluorescence resonance energy transfer (smFRET) with an *in vitro* eukaryotic translation system to investigate tRNA selection and subsequent steps during peptide elongation. We compared accommodation of a tryptophan-aminoacyl-tRNA into the ribosomal A-site containing either a cognate or near-cognate codon and unexpectedly found that, following an initial slow sampling event, subsequent near-cognate sampling events proceeded more rapidly than the initial event. Further, we found a strong negative correlation between the concentration of near-cognate aminoacyl-tRNA and the efficiency of tRNA accommodation. These novel characteristics of near-cognate interaction with the eukaryotic ribosome suggest that rejection of a near-cognate tRNAs leads to formation of an altered ribosomal conformation that assists in rejecting subsequent incorrect tRNA interactions.

## INTRODUCTION

Protein synthesis is a universally conserved process carried out by ribosomes, which are composed of ribosomal RNA (rRNA) and protein components organized into two subunits. The small subunit (“30S” in prokaryotes, “40S” in eukaryotes) is responsible for decoding the mRNA with the aid of transfer RNAs (tRNAs). The large subunit (“50S” in prokaryotes, “60S” in eukaryotes) is responsible for catalyzing the formation of peptide bonds between amino acids of the growing peptide chain (1–4). This process has been studied most extensively in bacteria.

The accurate decoding of the mRNA to produce a native peptide is essential to life and requires the efficient delivery of cognate aminoacyl tRNAs (aa-tRNAs), i.e., complementary to the 3-base mRNA codon, into the A-site of the ribosome. To facilitate this process, tRNAs charged with various amino acids are packaged into ternary complexes (TCs) with EF-Tu in bacteria (eEF1A in eukaryotes) and GTP which sample the A-site of the ribosome until a stable interaction between the anticodon stem loop of the cognate tRNA and the A-site codon is established. For cognate tRNA, codon recognition activates EF-Tu-catalyzed GTP hydrolysis and the departure of EF-Tu⋅GDP, thereby allowing the tRNA to accommodate into the A-site. The nascent peptide in the P-site is transferred to the new amino acid forming a new peptide bond and adding one amino acid to the nascent chain now linked to the A-site tRNA (5). Following this process, the two tRNAs are translocated from the A- and P-sites to the P- and E-sites, respectively, in a reaction catalyzed by the GTP complex of the translocase EF-G (eEF2⋅GTP in eukaryotes), and the elongation cycle is completed by dissociation of the deacylated tRNA from the E-site (1).

Initial binding among the 40 – 60 different TCs to the A-site of the ribosome is not codon sequence dependent, leading to the requirement of a powerful selection process that rejects incorrect (non-cognate and near-cognate) TCs that sample the A-site (6). Failure to exclude these tRNAs potentially leads to protein aggregation and loss of proteostasis in cells (7, 8). Consequently, ribosomes have mechanisms to provide exceptional fidelity during tRNA selection with an estimated error rate of approximately one in 10,000 codons translated in Saccharomyces cerevisiae (9, 10). Most of these errors are due to the misincorporation of near-cognate tRNAs, which have one incorrect base among the three anti-codon bases in the tRNA (9).

This high level of fidelity is established through kinetic steps during selection. While the stable 3-base Watson-Crick pairing interaction between the mRNA codon and the tRNA anti-codon triggers GTP hydrolysis and dissociation of EF-Tu⋅GDP, non-cognate aminoacyl tRNA rarely triggers GTP hydrolysis and instead dissociates promptly from the ribosome (1, 11, 12). Near-cognate tRNAs often foster GTP hydrolysis by ribosome-bound EF-Tu but are rejected in a subsequent proof-reading step, possibly due to the failure of the 30S subunit to undergo domain closure (13).

The process by which the ribosome discriminates between cognate and near-cognate tRNAs has been studied in depth in prokaryotes (5), but much less is known about the mechanistic details of this process in eukaryotes. Although the ribosome’s role in protein synthesis is preserved in all known forms of life, the overall composition of the ribosome varies significantly between Archaea, Bacteria, and Eukarya and even amongst the various species in each domain (14). These variations in the composition of the ribosome likely impact the dynamics and reaction pathways of protein synthesis. For example, in *E. coli* ∼20 amino acids are added to a nascent peptide per second during peptide elongation, whereas in higher eukaryotes the elongation rate is only ∼3 - 6 amino acids per second (15–17).

Kinetic processes are often compromises between speed and accuracy, so differences in elongation rate are likely to impact how the ribosome interacts with the repertoire of tRNA species in the cytoplasm (18, 19). Here we characterize the tRNA selection step of the peptide elongation cycle for cognate and near-cognate mRNAs in a reconstituted *in vitro* eukaryotic system, denoted PURE-Lite (28) using fluorescently labelled tRNAs and single molecule fluorescence resonance energy transfer (smFRET) experiments. We find that when ribosomes in the post-translocation (POST) state are exposed to near-cognate ternary complexes, the initial non-productive tRNA sampling event typically occurs much more slowly than subsequent rebinding events. Additionally, we find, unexpectedly, that the number of non-productive sampling events before successful accommodation increases significantly with TC concentration. We propose a kinetic model that closely predicts these results.

We also examined eEF2-catalyzed translocation of the mRNA and tRNAs in the A- and P-sites ribosomes containing either cognate or near-cognate aa-tRNA in the A-site. Our results with cognate peptidyl-tRNA are similar to those of others (21–23) except that, unlike that prior work, we find that tRNA dissociates from the E-site promptly following translocation, without requiring prior binding of the next TC into the A-site. Our results with near cognate tRNAs indicate that the translocation rate differs for stalled and active ribosomes.

## MATERIAL AND METHODS

### Preparation of Labeled tRNA

Bulk E. coli and yeast tRNA were purchased from Roche, Inc. Specific non-labelled tRNA isoacceptors were prepared from bulk *E. coli* tRNA (tRNA^Gln^, tRNA^Lys^) or bulk yeast tRNA (tRNA^Arg^, tRNA^Phe^, tRNA^Trp^, tRNA^Val^) via hybridization to bead-immobilized DNA oligos which were complementary to each specific tRNA isoacceptor and linked to the column beads by a 3’ biotin and streptavidin linkage, as described previously (24, 25). To minimize non-specific binding of labelled tRNAs to the PEG-passivated slides during single-molecule experiments, fluorescent labelled tRNA^Gln^ and tRNA^Trp^ were isolated instead using 5’ - CAA AAA CCG GTG CCT TAC CGC TTG GCG - 3’/3AmMO/ and 5’ - TTT GGA GTC GAA AGC TCT ACC ATT - 3’/3AmMO/ DNA oligos, respectively, with 3’ amino-modifications (3AmMO, IDT) conjugated to N-Hydroxysuccinimidyl (NHS)-Sepharose 4 Fast Flow beads (H8280, SIGMA).

To conjugate the DNA oligos, the NHS-Sepharose 4 Fast Flow beads were washed and pelleted extensively with 1 mM HCl. 500 nmol of amine derivatized oligo was added to 15 mL of packed bead slurry, mixed, and incubated for 2 hrs at room temperature followed by an overnight incubation at 4 °C on a rocking platform. Excess supernatant containing unreacted DNA oligo was removed by vacuum filtration. The beads were washed three times with 0.1 M Tris-HCl (pH 8.5, Buffer A), resuspended in Buffer A, incubated at room temperature for 4 hours to block unreacted succinimide, and then washed 4 times with Buffer A and 4 times with 0.1 M Na-acetate, 0.5 M NaCl (pH 5.0; Buffer B). Wash steps with Buffer A and Buffer B were repeated twice as described previously and beads were incubated overnight at 4 °C in Buffer B. The supernatant was then removed, and beads were washed with Buffer A repeatedly until the A_260_ value of the supernatant no longer decreased with subsequent washes. Using these amine-conjugated beads, tRNA^Gln^ and tRNA^Trp^ were then purified from bulk tRNA, reduced, and labelled with either Cy3- or Cy5-fluorescent dyes as described previously (24–27).

### Preparation of mRNA Sequences

All mRNAs used in these studies were transcribed from plasmids with the HiScribe T7 Quick High Yield RNA Synthesis Kit (E2050S, New England Biolabs). Transcripts contained a T7 promoter sequence (5’ – TAA TAC GAC TCA CTA TAG GG – 3’) followed by a sequence encoding the cricket paralysis virus internal ribosome entry site (IRES) which allows for initiation of the elongation cycle without the need for canonical initiation factors (28) (5’ - AGA CCG GAA TTC AAA GCA AAA ATG TGA TCT TGC TTG TAA ATA CAA TTT TGA GAG GTT AAT AAA TTA CAA GTA GTG CTA TTT TTG TAT TTA GGT TAG CTA TTT AGC TTT ACG TTC CAG GAT GCC TAG TGG CAG CCC CAC AAT ATC CAG GAA GCC CTC TCT GCG GTT TTT CAG ATT AGG TAG TCG AAA AAC CTA AGA AAT TTA CCT – 3’) and a coding sequence containing one of three codons at the sixth position (underlined) of the coding segment (28): **i**. a tryptophan codon (5’ - TTC AAA GTG AGA CAA <u>TGG</u> CTA ATG ACA TTT CAA GAT ACC – 3’); **ii**. a stop codon (5’ - TTC AAA GTG AGA CAA <u>TGA</u> CTA ATG ACA TTT CAA GAT ACC – 3’); or **iii**. a cysteine codon (5’ - TTC AAA GTG AGA CAA <u>TGC</u> CTA ATG ACA TTT CAA GAT ACC – 3’). This was followed by a partial Fluc+ gene sequence (5’ – ATG GAA GAC GCC AAA AAC ATA AAG AAA GGC CCG G – 3’) used as a spacer between the coding sequence and the 3’ end which terminated with biotin for attachment to the streptavidin-coated microscope slides used in single-molecule experiments.

Addition of a 3’-biotin modification to each mRNA was performed as previously described (29, 30).

### Preparation of Ribosome Complexes

40S and 60S ribosomal subunits were purified from shrimp (*Artemia salina*) cysts and eEF1A and eEF2 were purified from yeast (*Saccharomyces cerevisiae*), as described previously (31). Purified 40S and 60S ribosomal subunits were incubated with eEF1A, eEF2, Phe-tRNA^Phe^, Lys-tRNA^Lys^, Val-tRNA^Val^ and Arg-tRNA^Arg^, all at 2 mM, 2 mM GTP and the 3’-biotinylated mRNA IRES containing either a UGG codon (cognate to Trp-tRNA), a UGA codon (stop, near-cognate to Trp-tRNA), or a UGC codon (cysteine, near-cognate to Trp-tRNA) at the sixth position of the coding sequence. Post-translocation complexes positioned at the 4^th^ codon of the mRNA coding sequence (POST4 complexes) with a nascent peptide (FKVR) in the P-site were then formed (halted there by lack of the next ternary complex in the sequence, Gln-TC) and purified as described previously (30). Aliquots were then flash frozen in LN_2_ and stored at -80 °C.

### Preparation of PEG-passivated Flow Chambers

PEG-passivated slides were prepared as previously described, with some minor modifications (32). In brief, microscope slides (25 x 75 x 1 mm^3^, Globe Scientific, Inc) were drilled with a diamond-tipped bore producing two rows of 5 equally spaced holes for inflow and waste. Slides were washed thoroughly with deionized water and sonicated at 40 °C **i**. once for 25 min. in deionized water; **ii**. twice for 15 min. in acetone; and **iii**. once for 25 min. in methanol. They were then washed with deionized water, sonicated in 5 M KOH at 40 °C for 1 hour, washed 20 times with deionized water and three times with 200 proof EtOH. Slides were next dried using N_2_ gas and plasma cleaned at 800 - 1000 mTorr air pressure. They were then **i**. incubated overnight at room temperature in a silanization solution containing 1 ml 3-aminopropyltriethoxysilane, 5 ml acetic acid, and 94 ml methanol; **ii.** washed again 20 times with deionized water 20 times; **iii.** sonicated at room temperature for 10 minutes; **iv**. washed with EtOH 5 times; and v. dried with N_2_ gas. A polyethylene glycol solution (PEG, Laysan Bio, Inc., containing 20% (w/w) mPEG succinimidyl valerate, MW 2000 and 1% biotin-PEG-SC, MW 2000 dissolved in 0.1 M sodium bicarbonate (pH 8.3)) was then sandwiched between the salinized glass slide and a coverslip and incubated for at least 16 hours in a humidified chamber. Excess polyethylene glycol solution was then washed off with deionized water and the glass slides and coverslips were then immediately used to make 5-channel stopped-flow chambers with double-sided tape, as described previously (30), and stored in N_2_-purged Falcon tubes.

### Immobilization of Ribosome Complexes on PEG-passivated Slides

Immediately before the start of the experiment, 0.5 mg/mL streptavidin was injected into a stopped-flow chamber and incubated for 6 minutes. Unbound Streptavidin was washed out from the flow chamber with Buffer C (40 mM HEPES, 80 mM NH_4_Cl, 5 mM Mg(OAc)_2_, 100 mM KOAc, and 3 mM 2-Mercaptoethanol). For experiments measuring different aspects of tRNA selection and translocation in actively translating ribosomes, biotinylated POST5 complexes containing FKVRQ-Cy3-tRNA^Gln^ in the P-site of the ribosome were formed fresh at room temperature by incubating 0.01 µM POST4 complexes with 0.08 µM eEF1A, 1 µM eEF2, 2 mM GTP, and 0.03 µM Gln-tRNA^Gln^ (Cy3) for 6 minutes. Formation of POST5 complexes positioned the UGG codon (cognate to Trp-tRNA), UGA codon (stop, near-cognate to Trp-tRNA), or UGC codon (cysteine, near-cognate to Trp-tRNA) in the ribosomal A-site, according to the mRNA sequences described above. Alternatively, for experiments measuring translocation in stalled ribosomes, eEF1A, eEF2, and Gln-tRNA^Gln^ (Cy5) were incubated with POST4 complexes for 6 minutes to form biotinylated POST5 complexes containing FKVRQ-tRNA^Gln^ (Cy5) in the P-site. The biotinylated POST5 complexes were then injected into the streptavidin-coated flow chamber and incubated at 24°C for 6 minutes. Unbound ribosomes were washed out using Buffer C.

### Single-Molecule Fluorescence Imaging

Single molecule FRET studies were carried out at 24 °C. Unless otherwise stated, all fluorescence recordings were carried out in Buffer C supplemented with an enzymatic oxygen scavenging system of 2 mM protocatechuic acid (PCA), 50 nM protocatechunate 3,4 dioxygenase (PCD, Sigma Aldrich), 1 mM cyclooctatetraene (COT, Sigma Aldrich), 1 mM 4-nitrobenzyl alcohol (NBA, Sigma Aldrich; Aitken et al., 2008), and 1.5 mM 6-hydroxy-2,5,7,8-tetramethyl-chromane-2-carboxylic acid (Trolox, Sigma-Aldrich).

Image stacks were recorded at 25 s^-1^ frame rate on a custom-built objective-type total internal reflection fluorescence (TIRF) microscope (32) based on a commercial inverted microscope (Eclipse Ti-E, Nikon) with a 532-nm TIRF excitation laser. For measurements of tRNA selection, video recording was started and then various concentrations of Trp-tRNA^Trp^ (Cy5) in complex with eEF1A and GTP (Trp-TC (Cy5)) were injected under program control one second later. Co-injection experiments with Trp-tRNA^Trp^ (Cy5)·eEF1A⋅GTP and eEF2·GTP (Fig. S10 and S12) to study actively translating ribosomes were performed similarly using 25 nM of Trp-tRNA^Trp^ (Cy5)·eEF1A·GTP and 5 µM of eEF2·GTP. In experiments to study translocation in stalled ribosomes, Trp-tRNA^Trp^ (Cy3)·eEF1A·GTP was first injected into flow-chambers and incubated with Cy5-labeled POST5 complexes assembled previously for 6 minutes to form PRE6 complexes. Excess Trp-tRNA^Trp^ (Cy3)·eEF1A⋅GTP was then washed out of the flow-chamber and various concentrations of eEF2·GTP were injected 3 seconds after starting the recording. Measurements of non-specific (E-site) binding of uncharged tRNA^Trp^ (Cy5) to POST5 complexes containing Cy3-labeled glutamine tRNA in the P-site were done using 9.3 nM of uncharged tRNA^Trp^ (Cy5) (Fig. S10).

A deadtime of 0.7 seconds between start of the injection and arrival of fluorescence substrates at the sample surface was recorded for our injection system. This deadtime, plus the 1 or 3 s recorded time prior to injection, was subtracted from relevant measured waiting times (e.g. delay before Trp-TC (Cy5) binding). Single-molecule traces, unless otherwise stated, were filtered using a generalized Chung-Kennedy filter (33). Custom LabView scripts were used to trigger laser shutters and pumps.

Cumulative distributions of waiting times were fitted to single- or double-exponential functions using MEMLET (34). 95% confidence intervals were generated for these fits using bootstrapping with 500 iterations in MEMLET. *T*_Low Binding_, *τ*_Low Binding_, *T*_High Binding_, *τ*_High Binding_, *T*_Mid Binding_, and *τ*_Mid Binding_ for cognate and near-cognate conditions (Fig. 2 A; Table S1; Table S3) were measured manually by selecting the start and/or end times for each state. Likewise, *T*_SB-No Sampling_, *T*_SB-With Sampling_, and *T*_SB-All Traces_ (Table S2; Table S4) were measured manually by selecting the start times for stable binding events for cognate and near-cognate conditions. *N*_s_ was measured by counting the number of sampling events that occurred prior to stable binding by eye. Measurements of *T*_Subsequent Sampling_, *τ*_Initial_, and *τ*_Subsequent_ were done using the ebFRET analysis package (35). Hidden Markov Models were generated in ebFRET by fitting data with a 2-state model, with 10 restarts and precision set to 10^-8^.

Unless otherwise specified, uncertainties for all observed rate constants were determined using 95% confidence intervals. Uncertainties for FRET efficiency measurements (*E)* were calculated using standard deviation (s.d.) whereas the uncertainties for arrival time measurements (*T*) and lifetime measurements (*τ*) were calculated using standard error of the mean.

## RESULTS

To obtain insights into the molecular mechanisms of cognate and near-cognate tRNA selection using single-molecule FRET, we prepared eukaryotic ribosome POST complexes assembled on 3’-biotinylated mRNAs containing a pentapeptidyl-tRNA (FKVRQ-tRNA^Gln^ (Cy3)) in the P-site and either a cognate tryptophan codon (UGG) or a near-cognate stop codon (UGA) in the empty A-site (see Materials and Methods), as characterized previously by our group (20, 31). POST complexes containing the cognate tryptophan or near-cognate stop codon in the A-site are termed Trp-POST5 or Stop-POST5 complexes, respectively (Fig. 1 A and C).

**Figure 1:**
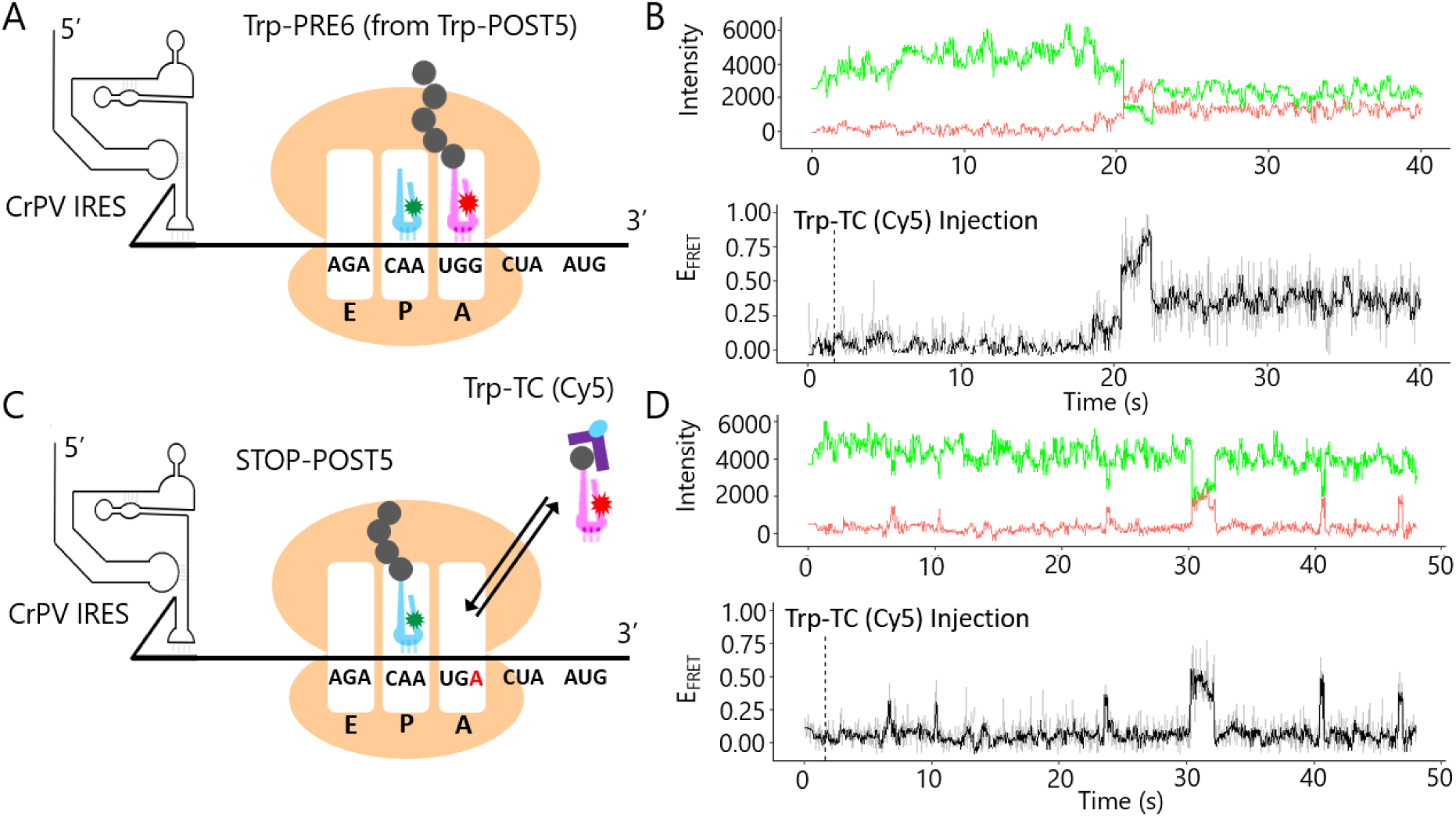
Schematic representations of ribosome complexes used in experiments and corresponding single-molecule traces. A). PRE6 ribosome resulting from Trp-TC binding to a POST5 complex assembled on an mRNA with a UGG tryptophan codon positioned in the A-site. B). Representative single-molecule fluorescence recording for cognate mRNA. Video recording was initiated, and Trp-TC (Cy5) was injected 1 s later (dashed vertical line). Green and red traces (top panels) show the Cy3 fluorescence emission from an individual ribosome and the Cy5 sensitized emission signal, respectively. Black traces (bottom panel) show the calculated FRET efficiency. Unfiltered data are shown in gray. Green, red, and black traces show data filtered using a generalized Chung-Kennedy Haran filter (33). C). POST5 ribosome with a UGA stop codon in the A-site. D). Representative single-molecule trace for near-cognate mRNA. Procedure and trace identities as in B.

**Figure 2:**
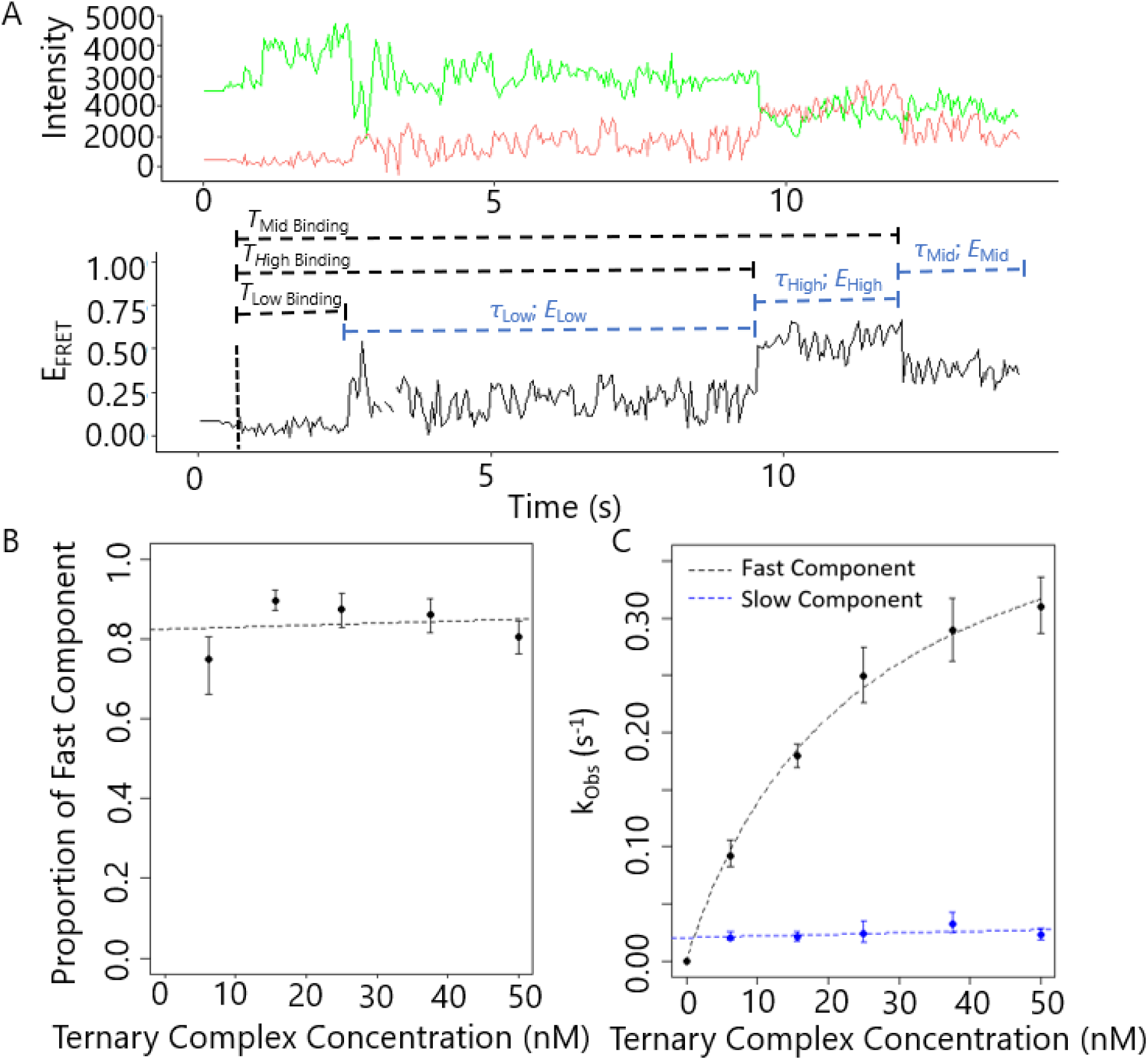
Kinetics of TC binding for cognate tRNA. A). Representative single molecule fluorescence trace showing timing parameters associated with stable binding of Trp-TC (Cy5) to the ribosome. Conditions and trace identities as in Fig. 1. B) and C). Kinetics obtained by maximum likelihood fitting with the sum of two exponential decays of the arrival times for Trp-TC (Cy5) FRET leading to stable binding. B). Relative amplitude of the faster exponential phase vs. TC concentration. C). Fast and slow fitted exponential rate constants. Error bars represent 95% confidence intervals generated by bootstrapping in MEMLET (34).

### Kinetics of Cognate tRNA Selection

Previous work has shown that aminoacyl-tRNA·eEF1A·GTP ternary complexes (TCs) often bind non-productively and dissociate before accommodation into the ribosome and peptidyl transfer can occur. This is common for non-cognate and near-cognate TCs and in some cases for cognate TCs as well (36, 37). To investigate the kinetics of cognate tRNA selection in our eukaryotic *in vitro* system, we immobilized Trp-POST5 complexes onto streptavidin-coated glass coverslips of stopped-flow cover-slip chambers and washed away unattached materials. Video recording of Cy3 and sensitized Cy5 fluorescence under TIRF illumination was initiated, after which cognate Trp-tRNA^Trp^ (Cy5)·eEF1A·GTP ternary complex (Trp-TC (Cy5)) was flowed into the chambers to initiate the tRNA selection process.

Binding of Trp-TC (Cy5) into the A-site of Trp-POST5 complexes gave rise to sensitized emission due to FRET between the Cy3- and Cy5-labeled tRNAs, and eventually resulted in accommodation and formation of long-lasting / stable PRE6 complexes, containing hexapeptidyl-tRNA FKVRQW-tRNA^Trp^ (Cy5) in the A-site and tRNA^Gln^ (Cy3) in the P-site (Fig. 1 A, B). FRET signals that resulted from Trp-TC (Cy5) binding events and persisted for at least 10 seconds prior to the disappearance of the FRET signal due to photobleaching were categorized as long-term accommodation events (stable binding events) and were found in the majority of single-molecule recordings in the cognate tRNA condition (∼82%; Fig. S2 A and D). The remaining traces either showed at least one non-productive Trp-TC (Cy5) binding/dissociation (sampling) event prior to accommodation (∼10%; Fig. S2 C and D) or failed to undergo productive accommodation (∼8%; Fig. S2 B and D), showing only brief non-productive sampling events prior to photobleaching of the tRNA^Gln^ (Cy3) in the P-site. Dissociation of tRNAs bound for >10 s followed by subsequent non-productive sampling events, indicative of tRNA proofreading, was extremely rare for cognate tRNAs (Fig. S1 A-C).

Approximately 60% of long-term accommodation events were immediately preceded by a combination of two transient FRET states (Fig. 2A). Transitions were common from near-zero background FRET to an initial low-efficiency FRET state of *E*_Low_ = 0.23 ± 0.10 (s.d., n = 539 traces; Fig. S3 A, B). The background or *E*_Low_ FRET state could also transition to a transient high-FRET state with a FRET efficiency of *E*_High_ = 0.49 ± 0.18 (s.d., n = 494 traces) and a mean lifetime τ_High_ = 4.97 ± 0.23 s (n = 960 traces; Fig. 2 A, Fig. S3 C and D). Similar intermediate FRET transitions have been found in previous studies of both bacterial and eukaryotic ribosomes (21, 38, 39). Differences between FRET efficiencies for these states measured here and in previous studies are most likely due to differences in fluorophore labeling positions on the tRNAs and differences in ribosome source. Based on these previous studies in prokaryotic and eukaryotic systems, it is likely that *E*_Low_ is associated with a codon decoding complex in which the Trp-TC (Cy5) interacts with the A-site mRNA, whereas *E*_High_ is a subsequent activated state occurring either before or after GTP hydrolysis and eEF1A-GDP dissociation (21, 37, 38) but prior to accommodation and peptide transfer. Following the formation of either *E*_Low_ or *E*_High_, the FRET efficiency stabilized at a medium FRET state with an average efficiency of *E*_Mid_ = 0.34 ± 0.16 and a lifetime limited by the photobleaching rate, again indicative of long-term accommodation.

The proportion of accommodated ribosomes that demonstrated the *E*_Low_ state, *E*_High_ state or both *E*_Low_ and *E*_High_ states prior to the formation of the accommodated *E*_Mid_ state (Fig. S4) did not vary significantly when tested from 6.2 nM to 50 nM Trp-TC (Cy5). The absence of detectable *E*_Low_ or *E*_High_ states from individual traces may be due to noise or to true differences between ribosomes measured in our experiments. FRET efficiency values (*E*_Low_ and *E*_High_) and their state lifetimes (τ_Low_ and τ_High_) are listed in Tables 1 and S1.

The time intervals for TC arrival leading to stable binding following TC injection, regardless of the initial FRET value (*E*_Low_, *E*_High_ or *E*_Mid_) or presence of sampling events prior to stable binding, exhibited fast and slow phases (Fig. 2 and S5 A) and were fit by double exponential functions over a range of Trp-TC (Cy5) concentrations (6.2 nM to 50 nM TC, Fig. S5 A). Maximum likelihood estimation (34) was used to obtain observed rate constants for Trp-TC (Cy5) arrival (Fig. 2 C). Approximately 80-90% of the recordings occupied the faster phase of stable binding time independent of the TC concentration (Fig. 2 B). The dependence on Trp-TC (Cy5) concentration of this faster component was well described by the Michaelis-Menten equation (Fig. 2 C), yielding *k*_cat_, *K*_m_, and *k*_cat_/*K*_m_ values given in Table 2. The saturation of the fast component at high TC concentrations suggests that a transient intermediate exists which does not give rise to a FRET signal prior to the first observed FRET increase. The slow component of the stable binding time distribution accounted for ∼10-20% of the recordings (Fig. 2 B) and its rate constant, 0.04 ± 0.01 s^-1^, was independent of TC concentration (Fig. 2 C). The time measurements used to obtain these parameters, as well as additional kinetic parameters, are given in Table S2.

### Kinetics of Near-Cognate tRNA Selection

In agreement with previous studies (5, 18), injection of Trp-TC (Cy5) near-cognate ternary complexes to Stop-POST5 complexes (Fig. 1 C) primarily led to transient and non-productive sampling events, with the majority (∼76%) of single-molecule traces at 9.3 nM Trp-TC (Cy5) not resulting in accommodation within the 300 s duration of the recordings (Figs. 1 D, S6 B and D). The remaining traces contained either both sampling events and stable binding (16%, Figs. S6 C and D) or only stable binding without prior sampling (8%, Figs. S6 A and D). The proportion of recordings with only sampling events decreased as Trp-TC (Cy5) concentration increased (Fig. S6 D), with a corresponding increase in stable complex formation. Approximately 40% of the ‘stable’ (≥10 s duration) complexes formed in the near-cognate case showed tRNA dissociation within the length of the recording and were followed by further Trp-TC (Cy5) sampling events, a much higher proportion than in the cognate case. This result suggests that tRNA proofreading is still possible following the formation of the *E*_Mid_ state (Fig. S1 D-F).

As for the cognate experiments, approximately 50% of the near-cognate stable binding events could be divided into distinct FRET states for *E*_Low_, *E*_High_, and *E*_Mid_, the values of which did not differ significantly from those measured with cognate tRNA (Table 1). Likewise, the proportion of stable binding events that contained *E*_Low_, *E*_High_, or both states prior to the formation of *E*_Mid_ were similar between cognate and near-cognate conditions (Fig. S4). However, durations *τ*_Low_ and *τ*_High_ for the near-cognate case were significantly shorter than for the cognate condition (Table 1; Table S3).

**Table 1:** States Formed on Cy5-TC Stable Binding to POST5 complexes. FRET efficiencies and durations for FRET states observed following stable binding of Trp-TC (Cy5) to POST5 complexes for cognate and near-cognate conditions.

| Table 1. States Formed on Cy5-TC Stable Binding to POST5 complexes |  |  |
| --- | --- | --- |
| Parameter | Cognate | Near-Cognate |
| $E_{Low}$ | $0.23 \pm 0.10$ (n = 539) | $0.2 \pm 0.1$ (n = 210) |
| $E_{High}$ | $0.5 \pm 0.2$ (n = 494) | $0.6 \pm 0.2$ (n = 193) |
| $\tau_{Low}$ | $4.4 \pm 0.2$ s (n = 1059) | $0.8 \pm 0.1$ s (n = 255) |
| $\tau_{High}$ | $5.0 \pm 0.2$ s (n = 960) | $2.2 \pm 0.2$ s (n = 199) |

**Table 2:**
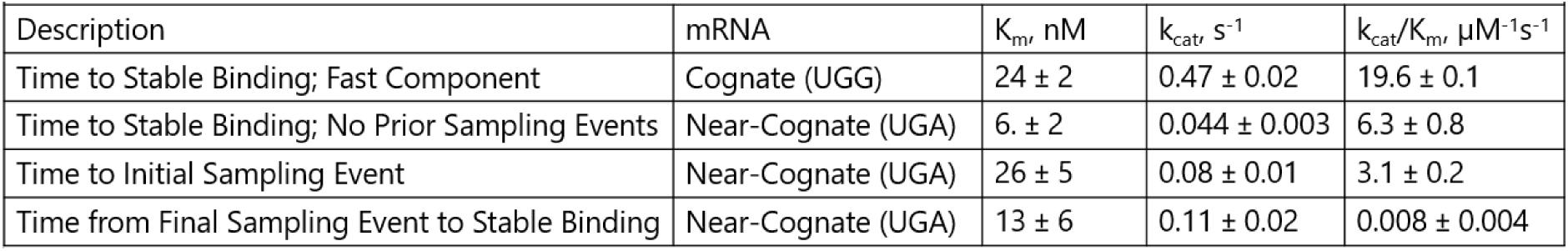
Binding Kinetics for Stable Binding and Sampling Events. Kinetic parameters associated with sampling and stable binding of Trp-TC (Cy5) to POST5 complexes for cognate and near-cognate conditions. Kinetic parameters were obtained by fitting Michaelis-Menten curves to observed rate constants for the various events in stable binding and sampling listed.

The times from exposure of Stop-POST5 complexes to Trp-TC (Cy5) to the initial non-productive sampling event (*T*_Initial Sampling_; Fig. 3 A) for the near-cognate case were fit by single-exponential functions to obtain observed rate constants which were well described by the Michaelis-Menten equation (Fig 3 B, D). This result is similar to the major component for the cognate case, but with a much slower saturating rate constant at high TC concentration (0.08 s^-1^ for the near-cognate interaction *vs.* 0.47 s^-1^ for the cognate interaction; Table 2). Because most near-cognate traces contained more than one non-productive sampling event prior to either the end of the trace or tRNA accommodation, we further measured the time between sampling events (*T*_Subsequent Sampling_; Fig. 3 A) that occurred following the initial non-productive event. Surprisingly, at each Trp-TC (Cy5) concentration tested, *T*_Subsequent Sampling_ was much shorter than *T*_Initial Sampling_ (Fig. 3 B, C), and generated rate constants that varied linearly with Trp-TC (Cy5) concentration, yielding a 2nd order rate constant of 4.4 µM^-1^s^-1^ (Fig. 3 D). In traces which contained both sampling events and stable binding, the time between the final sampling event and stable binding (*T*_Final Binding_) had values falling between *T*_Initial Sampling_ and *T*_Subsequent Sampling_ and also generated rate constants that followed Michaelis-Menten kinetics (Table 2; Table S4).

**Figure 3:**
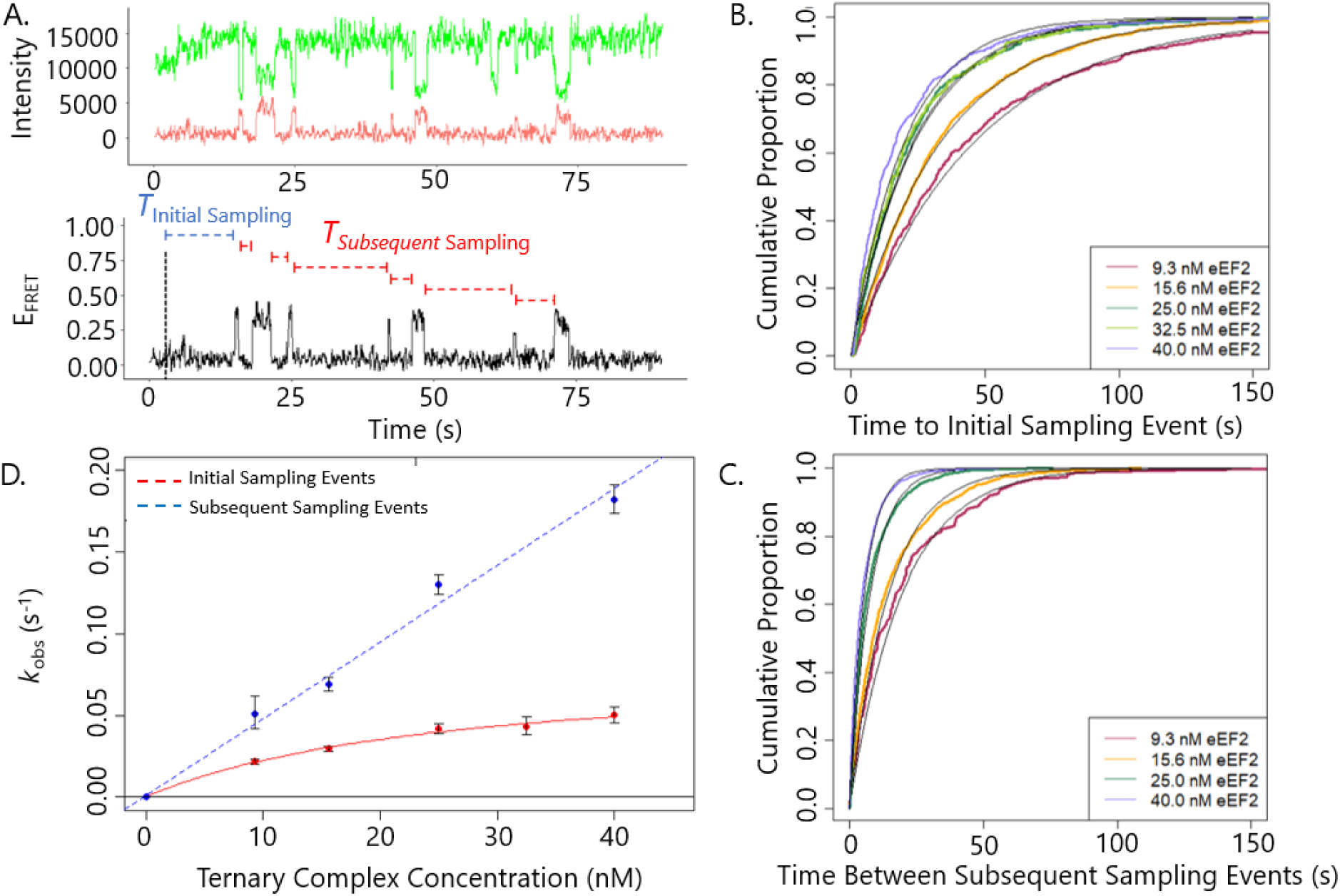
Kinetics of TC binding for near-cognate tRNA. A). Representative single-molecule trace and corresponding timing parameters of TC sampling the A-site. The thin vertical black line represents the time of TC injection. B). Cumulative distributions of the delay between delivery of Trp-TC (Cy5) and the initial A-site sampling event. Black lines represent single exponential fits. Each distribution represents at least 300 single-molecule recordings. C). Cumulative distributions of the delay between subsequent sampling events after the initial event. Each distribution represents ≥200 traces. D). Observed rate constants for initial and subsequent sampling events plotted versus TC concentration. Error bars represent 95% confidence intervals generated by bootstrapping.

The lifetimes of the initial and subsequent sampling events were indistinguishable from each other, giving values of 0.44 ± 0.06 s (n = 315 events) and 0.44 ± 0.03 s (n = 936 events), respectively, and demonstrated non-exponential lifetime distributions that suggest more than one kinetic step in series before tRNA dissociation (Fig. S7 A, B). The distribution of FRET efficiencies for these sampling events could be fit by the sum of two Gaussian components, having *E* values of 0.21 ± 0.08 and 0.48 ± 0.18 (n = 438 traces; 1383 sampling events; Fig. S8 A), which were close to the mean FRET efficiencies of the *E*_Low_ and *E*_High_ states for stable cognate and near-cognate binding.

A POST5 complex containing a different near-cognate codon (UGC, cognate to cysteine tRNA) in the A-site, was also found to exhibit a similar behaviour: binding of the TC for subsequent sampling events was faster than for initial non-productive events (Fig. S9 A). A more detailed study for this codon was not pursued due to the much lower number of stable binding events obtained relative to the UGA near-cognate stop codon (Fig. S9 B).

We considered three hypotheses which might explain this substantial increase in TC binding rate between the first sampling event following TC arrival and the subsequent ones. **1.** When solution is first flowed into the sample chamber it might not mix homogeneously with a surface layer where the ribosomes are located, so that the effective concentration of TC at the surface would be lower than at later times. **2.** The first TC to approach Stop-POST5 could bind to a testing (T-) site near the A-site or to one of multiple binding sites present on the ribosomal P-stalk of the large subunit, the position of which fluctuates near the mRNA/tRNA entry site of the ribosome (40). Thereafter a TC pre-bound to the ribosomal P-stalk might have preferential access to the A- or T-site, increasing the rate of binding, a sort of “channelling” of the substrates into the ribosome, as has been previously hypothesized (36, 40–42). **3.** When a near-cognate TC is rejected after a sampling event, the ribosome left behind is primed to bind further TCs faster due to a structural difference relative to the “naive” POST state that the ribosome assumes immediately following the previous elongation cycle.

We were able to test Hypothesis 1 by injecting 9.3 nM of uncharged Cy5-labeled tRNA^Trp^, which is expected to bind non-specifically to the E-site of Trp-POST5 complexes in a codon-independent manner to produce FRET with the Cy3-labeled P-site tRNA based on previous work in *E. coli* (43, 44). We found that the initial and subsequent binding events for the uncharged Cy5-labeled tRNA^Trp^ into the E-site were nearly identical to each other (Fig. S10 A). This, combined with the observation that cognate TC binding is also faster than the initial near-cognate interaction (compare *T*_Initial Sampling_ in Tables S1 and S3), indicates that the rates are not markedly compromised by inhomogeneous TC concentration (i.e. lower at the coverslip surface) early after injection, apparently ruling out Hypothesis 1.

Hypothesis 2 was tested by pre-incubating the Stop-POST5 complex with 5 µM eEF2·GTP to occupy the G-protein factor binding sites on the ribosomal P-stalk, thus preventing TC “channelling” in the subsequent step. When TC was then added in the presence of 5 µM eEF2·GTP, the initial and subsequent rates of TC binding to Stop-POST5 ribosome were similar to those without eEF2 present (Fig. S10 B), providing strong evidence against the channelling hypothesis.

Hypothesis 3, that “memory” of a near-cognate sampling event changes the kinetic properties of the ribosome and additional features of near-cognate proofreading is taken up in the Discussion.

### Cognate tRNA exhibits faster subsequent TC binding when accommodation is blocked

Tigecycline is an antibiotic that blocks aa-tRNA accommodation within the A-site of prokaryotic ribosomes without inhibiting EF-Tu-induced GTP hydrolysis and has also been shown to block accommodation in eukaryotic ribosomes (45). We reasoned that if we used tigecycline to inhibit the accommodation of the cognate tRNA, the dynamics of cognate tRNA sampling might mimic those of near-cognate tRNA sampling. Although sampling events with rejection of tRNA did occur with the cognate tRNA more often in the presence of 100 µM tigecycline, stable binding of tRNA was also common (data not shown). *T*_Initial Sampling_ and *T*_Subsequent Sampling_ (Fig. S11 A, B) arrival times were fitted by single exponential functions giving observed rate constants which again indicated that subsequent binding was faster than initial binding (Fig. S11 C), although the difference was smaller than with the near-cognate codon in the absence of the antibiotic. We observed stronger similarities between cognate tRNA in the presence of tigecycline and near cognate tRNA for **i.** the lifetimes of the sampling events, *τ*_Initial_ and, *τ*_Subsequent_, (Fig. S7 C, D), **ii.** the non-exponential lifetime distributions, again suggesting more than one kinetic step in series before the tRNA dissociates, and **iii.** the FRET efficiency distributions for the sampling events, with components at *E* = 0.21 ± 0.08 and *E* = 0.48 ± 0.18 (s.d., n = 393 traces; 2121 sampling events; Fig. S8 B). However, the *E*_High_ high-FRET state was more abundant for the tigecycline-induced sampling events than for near-cognate sampling events.

Assuming that tigecycline blocks aa-tRNA accommodation without inhibiting GTP hydrolysis with eukaryotic ribosomes as in prokaryotes, our results lead to two suggestions: 1. In the presence of the drug, more cognate TCs proceed through to GTP hydrolysis prior to rejection compared to near-cognate TCs in the absence of tigecycline. 2. More importantly, when accommodation is inhibited, the kinetic mechanism which results in subsequent sampling events proceeding more rapidly than the initial sampling events applies to both cognate and near-cognate tRNAs.

### Dependence of near-cognate accommodation on TC concentration

Due to the low-probability of accommodation of near-cognate tRNAs into the ribosome, single-molecule studies of near-cognate accommodation are mostly limited to the initial binding step (21). To compare the kinetics of cognate and near-cognate tRNA selection further, we relied upon a misincorporation-prone mRNA sequence for our near-cognate experiments (see Discussion). This allowed ∼20% of our near-cognate ribosomes to undergo accommodation within the length of our recordings without undergoing further rejection due to proofreading.

For the near-cognate condition, when the first binding event resulted in stable binding, the arrival times were similar to the time to the initial sampling event at various TC concentrations (Table 2, Table S4). In contrast, traces that contained non-productive sampling events prior to stable binding were far slower and, when fit to single-exponential distributions, generated rate constants that were similar (about half the rate) to the slow component of stable binding for the cognate case and did not vary appreciably with Trp-TC (Cy5) concentration (Fig. 4 D, E; Table S4).

**Figure 4:**
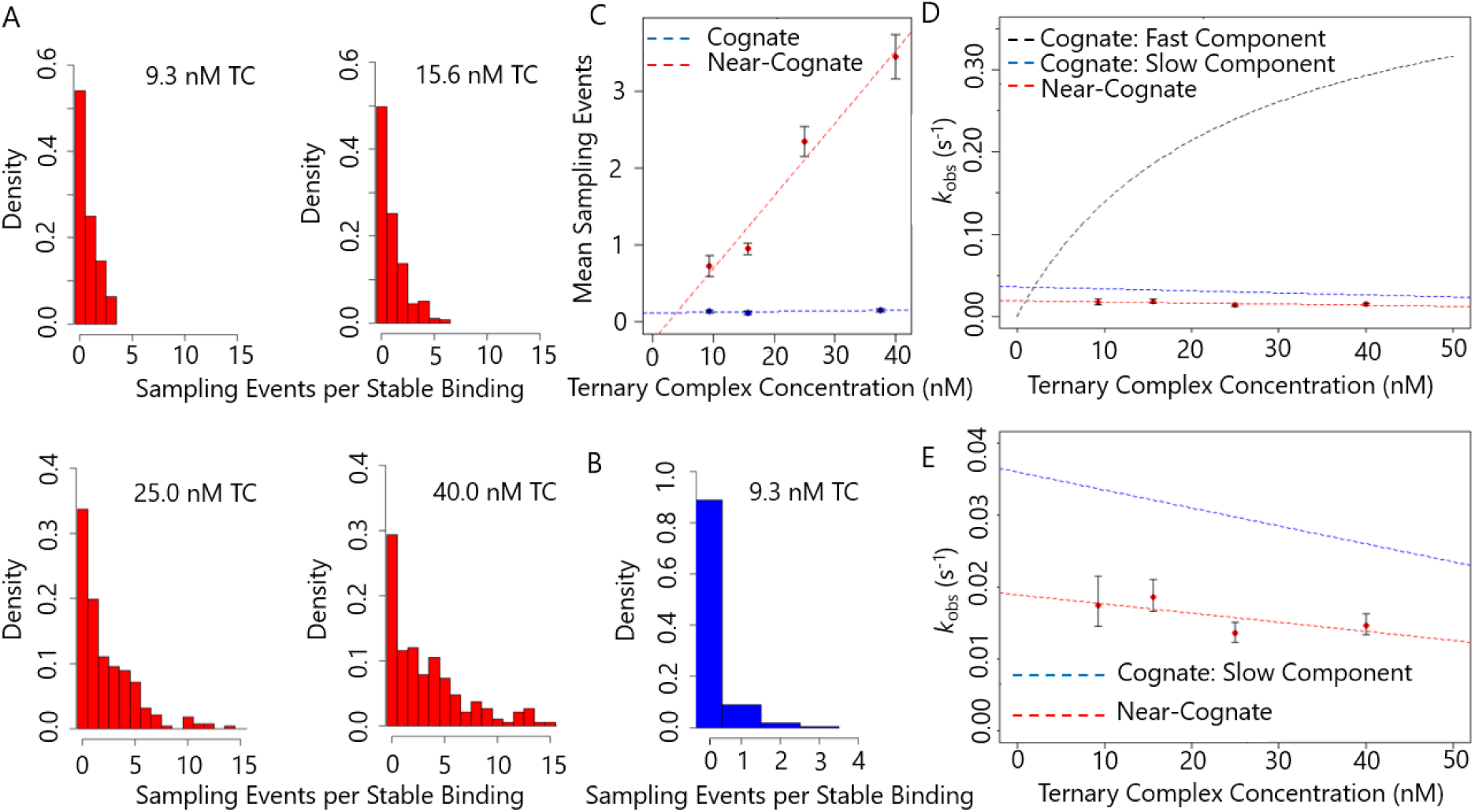
Kinetics of stable binding for near-cognate tRNA. A). Histograms representing the number of sampling events (with dissociation) that occur prior to stable binding for the near-cognate condition at listed TC concentrations. B). Representative histogram for the number of sampling events prior to stable binding for the cognate condition at 9.3 nM TC. The other TC concentrations gave the same result. C). Average number of sampling events prior to stable binding (N_S_) for the cognate (blue) and near-cognate (red) mRNA conditions. Error bars represent the standard error of the mean. Measurements for the near-cognate tRNA condition were generated from ≥ 200 traces, except for the 9.3 nM TC condition which was generated from ∼50 traces. Measurements for the cognate tRNA condition were generated from ≥ 200 traces. The line for cognate mRNA indicates the average among the TC concentrations. The line for the near-cognate mRNA (red) represents a linear regression fit. D). Observed rate constants (red) for the time from TC delivery to stable binding (k_SB-All Traces_) for the near-cognate tRNA condition. The fitted curves from Fig. 2C for the fast (black) and slow (blue) components of k_SB-All Traces_ for the cognate mRNA are shown for comparison. E). Vertically expanded portion of data from (D). Error bars for (D) and (E) represent 95% confidence intervals generated by bootstrapping.

Assuming the simplest hypothetical scheme for tRNA selection, in which each bound cognate or near-cognate tRNA has a fixed probability of accommodating into the ribosome *vs*. being rejected during tRNA selection (see Discussion), the number of sampling events that occur before stable binding should be independent of TC concentration. To test this hypothesis, we further determined the mean number of near-cognate sampling events that occurred prior to accommodation and, surprisingly, found that the mean number of sampling events before stable binding (*N*_s_) for the near-cognate tRNA varied strongly with Trp-TC (Cy5) concentration (Fig. 4 A and C). In contrast, *N*_s_ for the cognate tRNA was low (∼0.1 sampling event per stable binding event) and did not vary with Trp-TC (Cy5) concentration, as originally expected (Fig. 4 B, C). These results indicate that as the concentration of near-cognate Trp-TC (Cy5) is increased, the likelihood of accommodation following a sampling event decreases. A kinetic mechanism accounting for this observation is presented in the Discussion.

### tRNA Dynamics in PRE6 Complexes

Trp-PRE6 complexes with tRNA^Gln^ (Cy5) in the P-site and FKVRQW-tRNA^Trp^ (Cy3) in the A-site, stalled by the absence of eEF2, oscillated between a low-FRET and a higher-FRET state (Fig. S12 A), consistent with previous observations of tRNA fluctuations in stalled *E. coli* PRE complexes (46–48). Structural and single-molecule studies with cognate PRE complexes formed on *E. coli* ribosomes attributed the low-FRET state to tRNAs positioned in the classical A/A and P/P positions and the high-FRET state to tRNAs in the A/P and P/E hybrid positions and with the small ribosomal subunit rotated relative to the large subunit (32, 38, 49). The low-FRET state detected here had a mean lifetime of *τ*_Classic_ = 2.1 ± 0.1 s and a mean FRET efficiency of *E_Classic_* = 0.30 ± 0.11 (s.d., n = 535 traces; 2284 FRET transitions), whereas the high-FRET state had a mean lifetime of *τ*_Hybrid_ = 0.90 ± 0.04 s and a mean *E*_Hybrid_ = 0.45 ± 0.12 (s.d., n = 535 traces; 2175 events). Similarly, Stop-PRE6 complexes oscillated between a low-FRET state with *E*_Classic_ = 0.28 ± 0.12 (s.d., n = 142 traces; 2019 events) and *τ*_Classic_ = 1.09 ± 0.03 s and a high-FRET state with *E*_Hybrid_ = 0.36 ± 0.13 (s.d., n = 142 traces; 2019 events) and *τ*_Hybrid_ = 0.70 ± 0.01 s (Fig. S12 B, C). Our measurements agree roughly with values previously determined in similar studies using stalled *E. coli* ribosomes: *E*_Classic_ = ∼0.3, *τ*_Classic_ = ∼2 s, *E*_Hybrid_ = ∼0.65 and *τ*_Hybrid_ = ∼3 s (32, 47).

### eEF2-Catalyzed Translocation for Cognate and Near-Cognate mRNA

We compared the dynamics of eEF2-dependent translocation of biochemically stalled Trp-PRE6 and Stop-PRE6 complexes prepared by pre-incubating Trp-POST5 and Stop-POST5 complexes containing FKVRQ-tRNA^Gln^(Cy5) in the P-site with Trp-TC (Cy3) for ∼8 min prior to injection of eEF2⋅GTP. A representative trace of the time-dependent FRET changes obtained for translocation of Trp-PRE6 is shown in Fig. S13A. Prior to eEF2 injection, the Trp-PRE6 exhibited an equilibrated mixture of the *E*_Classic_ and *E*_Hybrid_ FRET states with an overall average FRET efficiency of *E*_Mid_ = 0.34 ± 0.16 (s.d., n = 497 traces, Fig. S13 B). Injection of eEF2⋅GTP into the stalled Trp-PRE6 sample led to a transition to a high-FRET state (*E*_trans_ = 0.59 ± 0.04, s.d., n = 1154) followed by complete loss of the FRET signal (Fig. S13 A, B). Transitions from *E*_Mid_ to *E*_trans_ showed a slight bias for the *E*_Hybrid_ state, with ∼70% of traces occupying the *E*_Hybrid_ state immediately prior to the transition to the *E*_trans_ state. The increase in the FRET efficiency is consistent with structural studies (50–53) showing that the P-and E-site (POST) tRNAs are more closely spaced in the ribosome than A-and P-site tRNAs, and as previously noted in single molecule experiments examining translocation in human ribosomes (23, 51). The apparent rate constant for formation of high FRET state increased as a function of eEF2 concentration, reaching a value of *k*_trans_ = 0.68 ± 0.06 s^-1^ (Fig. S13 C) at the nearly saturating concentration of 5 µM eEF2 (20). The subsequent loss of FRET is due to either dissociation of the E-site tRNA or photobleaching. Determining the rate constants for the duration of the high-FRET state at two laser intensities and at two near-saturating eEF2 concentrations (Fig. S14 A) allowed us to compensate for photobleaching and estimate the rate constant for E-site tRNA dissociation as *k*_diss_ = 0.67 ± 0.02 s^-1^ at 5 mM Mg^2+^, a value which showed little dependence on eEF2 concentration (Fig. S13 D), but decreased ∼3-fold when Mg^2+^ was raised to 15 mM Mg^2+^ (data not shown). The rate constant for E-site tRNA dissociation at 5 mM Mg^2+^ is ∼30-fold higher than earlier reported for human ribosomes (23), in which an initiation complex, prepared nonenzymatically using 15 mM Mg^2+^, was used for the elongation cycle. This large difference may be attributable to conformational changes induced by high Mg^2+^ concentrations which are not readily reversible when Mg^2+^ concentration is reduced.

Although there was a large reduction in the eEF2-dependent conversion of Stop-PRE6 to Stop-POST6 (30%) compared with conversion of Trp-PRE6 to Trp-POST6 (60%), similar results for Stop-PRE6 compared with Trp-PRE6 were obtained for the transient high FRET state (*E*_trans_ = 0.55 ± 0.20), which was formed with a rate constant of *k*_trans_ = 0.78 ± 0.03 s^-1^ at 5 µM eEF2, and for the rate constant for tRNA dissociation (*k*_diss_ = 0.41 ± 0.05 s^-1^). The similarity for the rate constants was unexpected, since ensemble studies^Ng21^ of the conversion of Stop-POST5 to Stop-POST6 on simultaneous rapid mixing with Trp-TC and eEF2·GTP showed translocation to proceed 20-fold more slowly than during similar conversion of Trp-POST5 to Trp-POST6, slow near-cognate tRNA translocation was also found in an *E. coli in vitro* reconstituted system (54). Our attempts to determine the dynamics of coupled Stop-PRE6 and Stop-POST6 formation by smFRET failed due to very low yields of Stop-POST6 formation (≤ 10% of Stop-POST5) on simultaneous addition of Trp-TC and eEF2 to Stop-POST5.

Earlier we described functional differences between stalled and actively elongating E. coli ribosomes (47) We speculate that the difference in translocation rates of Stop-PRE6 which we observe may be attributable, at least in part, to the smFRET studies being conducted on preformed PRE complexes, i.e., stalled ribosomes, whereas the earlier ensemble experiments measuring Stop-POST5 conversion to POST6 complexes proceeded via simultaneous addition of Trp-TC and eEF2, i.e. translocation by active ribosomes. Further experiments will be required to resolve this issue.

## DISCUSSION

Here we use single-molecule FRET to determine the kinetics of cognate and near-cognate tRNA selection and translocation in a reconstituted *in vitro* eukaryotic translation system with ribosomes programmed with CrPV-mRNA. Our results related to the kinetics of cognate and near-cognate tRNA selection and translocation are largely consistent with prior studies conducted in our labs on peptide elongation, as shown in Table S5 (20, 21) . As described above, however, for both cognate and near-cognate codons in ribosomes stalled by delay in adding eEF2·GTPP, translocation was prompt, whereas in our earlier ensemble studies when TC and eEF2 were added simultaneously, the near-cognate PRE complex translated slower than the cognate complex. Preliminary single molecule data suggest that the difference is due to stalling elongation vs. actively cycling ribosomes as reported before in E. Coli ribosomes (47). Additionally, our results describing near-cognate tRNA selection prior to accommodation are inconsistent with common models of tRNA selection that assume that non-productive dissociation of near-cognate aa-tRNA preceding peptide bond formation (i.e., the result of proofreading) results in simple reformation of the initial POST complex.

Specifically, the following three observations are incompatible with proofreading directly resulting in the reformation of an identical initial POST complex following near-cognate tRNA rejection. First, subsequent non-productive sampling events occur more rapidly compared to the initial non-productive sampling event following exposure of ribosomes in the post-translocation state to near-cognate TCs. Second, the average number of non-productive sampling events that occur before a near-cognate tRNA is successfully accommodated into the ribosome increases with near-cognate TC concentration (Fig. 3 B-D). Third, the overall rate of stable binding for the near-cognate tRNA does not vary significantly with the concentration of TC, despite faster TC binding at higher TC concentrations. These observations were made with two different near-cognate codons in the ribosomal A-site, (UGA, stop) and (UGC, Cys), and with the cognate codon (UGG, Trp) when accommodation was blocked by tigecycline. Taken together, they suggest that as the concentration of near-cognate TC is increased, an inhibitory mechanism related to incorrect base pairing decreases the likelihood of stable binding, and lead us to propose the new proofreading scheme, shown in Fig. 5 A (Scheme 1).

**Figure 5:**
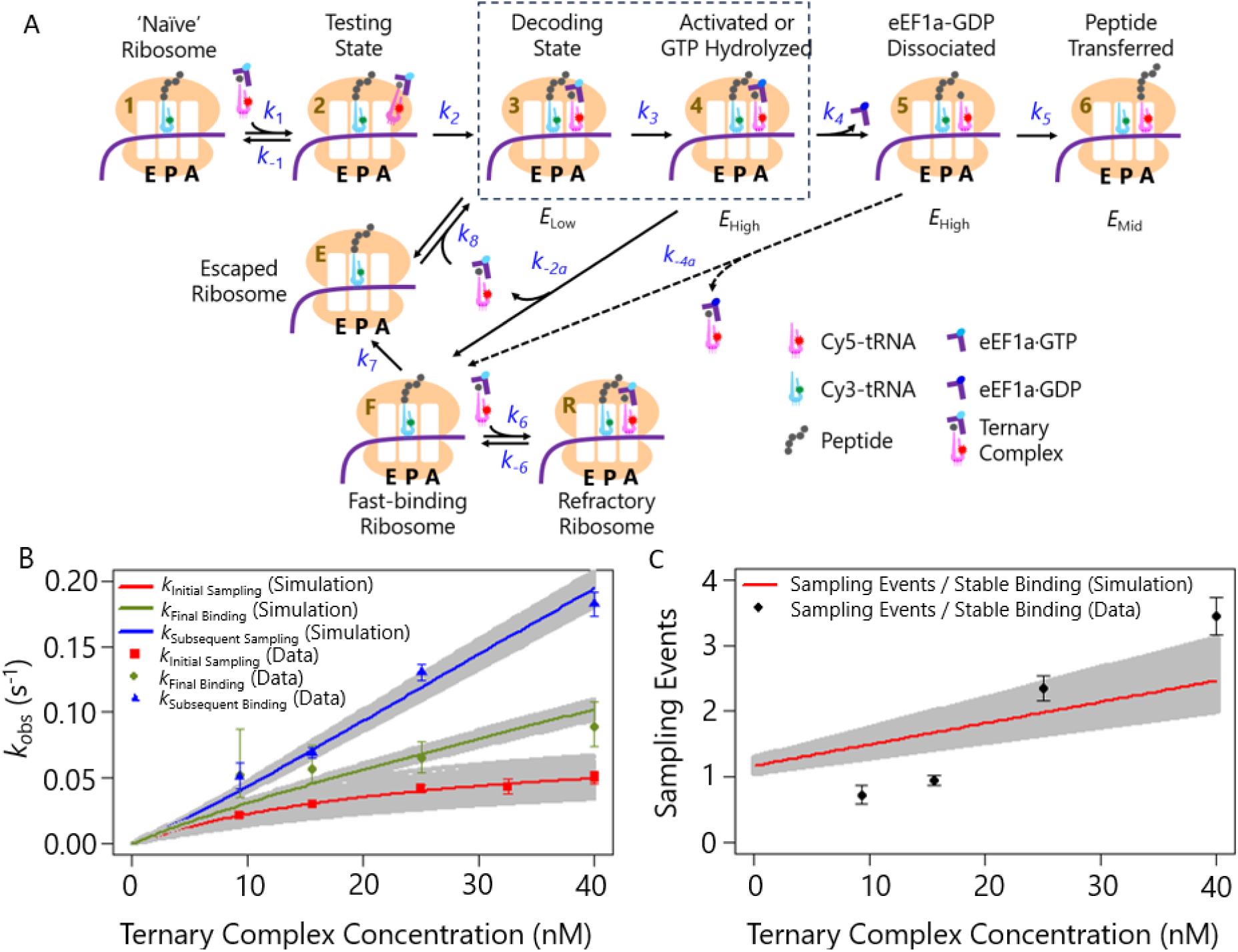
Kinetic reaction scheme that predicts the experimental results. A). The top line of the scheme represents a standard or typical model of tRNA selection (States 1 – 6). The present experimental results imply that ribosomes after dissociation of a near-cognate TC retain a structural conformation (State F) which binds TCs faster than State 1 and, following TC binding (State R), exhibits inhibition of accommodation relative to States 3 and 4. States 3 and 4 are considered together. Step 4b is dashed because it is uncertain and not required to explain the results. See text for more explanation. B). Fits from the analytical model (Supplementary Appendix A) to the observed rate constants for initial sampling events (Data: red squares; Model: red line), subsequent sampling events (Data: blue triangles; Model: blue line) and final sampling events prior to stable binding (Data: green circles; Model: green line). C). Model fit (red line) to the mean number of sampling events prior to stable binding at each TC concentration (black circles). Gray shaded regions around each model fit represent 95% confidence intervals determined from analytical model.

In Scheme 1, the initial POST complex, State 1, binds a near-cognate TC in two steps. Following arrival of the TC into the flow chamber, the rate of appearance of the first FRET-producing interaction (*E*_Low_) follows a hyperbolic relationship *vs*. TC concentration with *K*_1/2_ = 23 nM and a saturating rate constant of 0.08 s^-1^. The cumulative distribution of the arrival times (Fig. 2B) shows no lag phase, even at low TC concentrations, indicating that step 1 (States 1 ↔ 2) is a rapidly equilibrating reversible collision with *k*_-1_ >> *k*_2_. State 2 could be a “testing site” in which the TC is not making full contact with the mRNA (13) or it could correspond to binding at the P-stalk (40), neither of which comes close enough to transfer excited-state energy to the P-site tRNA (FRET *E* = 0). State 3 is the first intermediate which produces FRET (*E*_Low_) and is converted to State 4 relatively rapidly. *k*_4_ and *k*_5_ are rate constants for eEF1A dissociation, accommodation, and peptide transfer, respectively.

The accommodation of a near-cognate tRNA via steps *k*_3_ and onward would result in a misincorporation event that is typically prevented by the ribosome’s proofreading activity. The more likely event for a near-cognate TC is to non-productively sample the ribosomal A-site and dissociate from either State 3 or State 4 prior to peptide bond formation. Because States 3 and 4 are indistinguishable from each other with the current experimental results, we have combined them (dashed box) for the purposes of modelling the overall reaction scheme. According to the simplest model of tRNA selection, once a tRNA fails to undergo accommodation and dissociates from the ribosome, the ribosome returns to the same state that it was initially when it originally encountered the tRNA. However, because subsequent non-productive sampling events bind to the ribosome more rapidly than the initial sampling events, it suggests that the ribosome assumes an altered state following near-cognate tRNA rejection. Therefore, ribosomes in States 3/4 following tRNA rejection do not simply return to State 1 but instead proceed so an altered state, State F, via a reaction step with rate constant of *k*_-2a_. State F then either binds a near-cognate TC again (Step 6) to form State R, a FRET-detectible complex, or converts in a first order reaction to State E (Step 7). We posit that State R is refractory, or at least slow, towards undergoing GTPase activation or tRNA accommodation, necessary for peptide elongation, thereby delaying the ribosome in functionally processing a near-cognate TC. This echoes other results with near-cognate TCs or with inhibitory codon pairs which have provided evidence for persistent and essentially irreversible changes limiting the stoichiometry of peptide elongation (20, 55). Reversible near-cognate TC binding to form State R thus results in repeated unsuccessful sampling events. In competition with step 6, step 7 forms State E which binds near-cognate TC (Step 8) to reform State 3, providing a renewed opportunity to activate GTP hydrolysis, fully accommodate aa-tRNA within the A-site, and form the next peptide bond finally inserting the near-cognate amino acid into the peptide chain. We considered whether State F could convert directly to State 1, obviating the need to invoke State E, but rejected this possibility because we find the final transition to stable binding of near cognate TC after sampling events to proceed more rapidly than the initial transition (Fig. 5 B).

In Scheme 1, the relative rates of near-cognate TC binding resulting in State conversion falls in the order: State F to State R (*k*_6_) > State E to State 3 (*k*_8_) > State 1 to State 3 (*k*_1_·*k*_2_/*k*_-1_). This order of reactivity accounts for the observations that subsequent sampling events proceed more rapidly than the initial sampling event, and that this difference increases with increasing near-cognate TC concentration, which favors State F partitioning toward State R over State E. It also is consistent with our observation that the average number of near-cognate sampling events occurring before stable binding increases with TC concentration, reaching a value of 3.5 ± 0.3 sampling events per stable binding event at 40 nM TC (Fig. 5 C).

The plausibility of Scheme 1 in accounting for our observations is demonstrated by our ability to quantitatively fit the near-cognate kinetic data of Figs. 3 D, 4 B and 4 C to Scheme 1 using maximum likelihood optimization (Supplementary Appendix A). This permits estimation of some of the rate constants in Scheme 1, which are presented in Table 4 along with those determined directly from experimental data (see Materials and Methods). With these values, Scheme 1 predicts *k*_Initial Sampling_, *k*_Subsequent Sampling_, *k*_Final Binding_, the durations of the observed FRET states, and number of sampling events, *N*_s_, quite well (Fig. 5 B and C). Parameter confidence limits (gray bands) were determined by bootstrapping. We also programmed a Monte Carlo simulation of Scheme 1 which confirmed the analytical equations (data not shown).

**Table 3:** Translocation Kinetics. Kinetic parameters and FRET states associated with eEF2-catalyzed translocation from PRE6 to POST6 for cognate and near-cognate conditions. Kinetic parameters were obtained by fitting Michaelis-Menten curves to observed rate constants shown in Fig. S13.

| Table 3. Translocation Kinetics |  |  |
| --- | --- | --- |
| Parameter | Cognate | Near-Cognate |
| $E_{\text{trans}}$ | $0.59 \pm 0.04$, n = 1154 | $0.6 \pm 0.2$, n = 671 |
| $k_{\text{trans}}, \text{s}^{-1}$ | $0.8 \pm 0.1$ | $0.8 \pm 0.1$ |
| $K_{\text{m}}^{\text{eEF2}}, \mu\text{M}$ | $0.2 \pm 0.2$ | $0.3 \pm 0.1$ |
| $k_{\text{diss}}, \text{s}^{-1}$ | $0.67 \pm 0.02$ | $0.41 \pm 0.05$ |

**Table 4:**
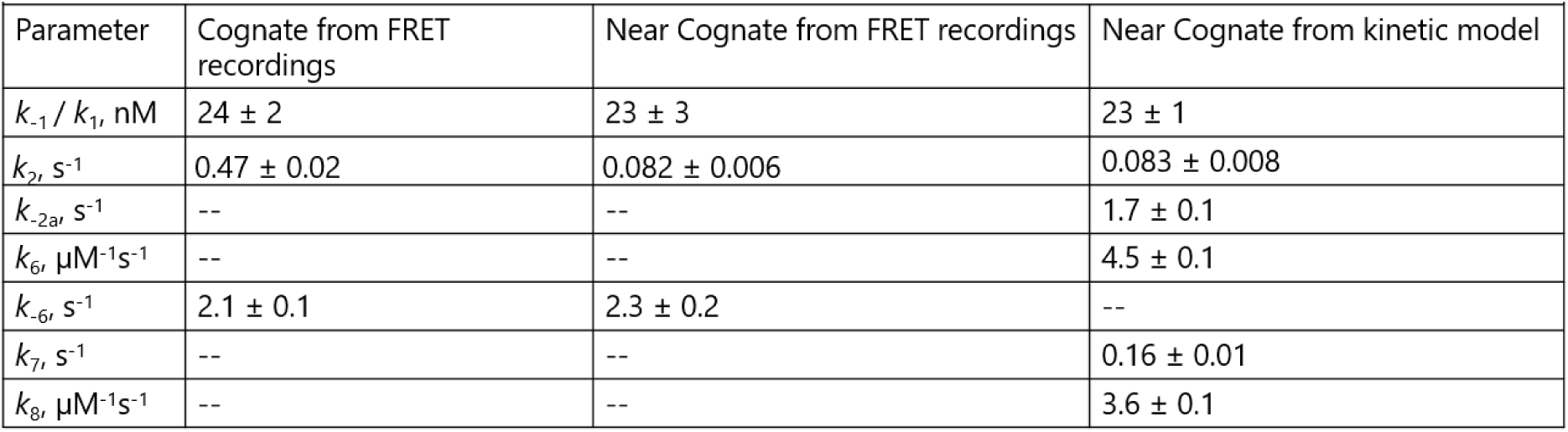
Rate constants for Cognate and Near Cognate recognition. Estimated rate constants of tRNA selection for cognate and near-cognate mRNA. Cognate rates determined from lifetimes of the FRET states, as in Figs. S3 and S7. Near cognate rates were determined either by measuring FRET state lifetimes (Fig. S3 and S7) or by fitting the kinetic scheme of Fig. 5 to the distributions of event durations and numbers of sampling events as explained in Supplementary Appendix A.

| Table 4: Rate Constants for Cognate and Near Cognate recognition |  |  |  |
| --- | --- | --- | --- |
| Parameter | Cognate from FRET recordings | Near Cognate from FRET recordings | Near Cognate from kinetic model |
| $k_{-1} / k_1$, nM | $24 \pm 2$ | $23 \pm 3$ | $23 \pm 1$ |
| $k_2$, $s^{-1}$ | $0.47 \pm 0.02$ | $0.082 \pm 0.006$ | $0.083 \pm 0.008$ |
| $k_{-2a}$, $s^{-1}$ | -- | -- | $1.7 \pm 0.1$ |
| $k_6$, $\mu M^{-1}s^{-1}$ | -- | -- | $4.5 \pm 0.1$ |
| $k_{-6}$, $s^{-1}$ | $2.1 \pm 0.1$ | $2.3 \pm 0.2$ | -- |
| $k_7$, $s^{-1}$ | -- | -- | $0.16 \pm 0.01$ |
| $k_8$, $\mu M^{-1}s^{-1}$ | -- | -- | $3.6 \pm 0.1$ |

The great majority of tRNAs in a cell are non-cognate or near-cognate to any given codon, so the fast-TC-binding binding conformation in our model (State F) is expected to be the predominant post-translocation intermediate for ribosomes containing only one tRNA, in accord with cryo-EM images of ribosomes in fast-frozen cells (56) showing a unique post-translocation state containing one tRNA. This is presumably State F, rather than State 1. The apparent inhibition of accommodation in State R, indicated by an increased number of sampling events that occur prior to stable binding as near-cognate TC concentration is increased, likely facilitates the rejection of near-cognate and, potentially, non-cognate TCs. The durations of the first sampling event and subsequent ones are the same indicating that the near-cognate TC rejection rate from state R (*k*_-6_) is approximately the same as from state 3/4 (k_-2a_). This equivalence leads to normal exchange between bound and free TCs in states F and R. The present data don’t provide information on whether State R would activate GTP hydrolysis and accommodation for cognate TCs (State R -> State 5, which we did not include for the near-cognate pathway). If further reactions from State R are not inhibited for cognate TCs, then this novel mechanism could contribute prominently to fidelity of tRNA selection in eukaryotic elongation by facilitating accommodation of cognate tRNAs.

## DATA AVAILABILITY

*The data underlying this article are available in* the Figshare repository, at https://dx.doi.org/[ 10.6084/m9.figshare.24581943].

## AUTHOR CONTRIBUTIONS

C.F., Y.E.G, and B.S.C. conceived and planned the experiments. C.F. carried out the experiments. C.F. analysed data generated from experiments. Y.E.G. and P.S.N. planned and carried out kinetic simulations presented in the work. C.F., A.B., M.Y.N., and H.L. contributed to sample preparation. C.F. wrote the manuscript. C.F., Y.E.G., B.S.C., and P.S.N. edited the manuscript.

## ACKNOWLEDGEMENTS

We thank Xiaonan Cui for assistance with protein purification and for additional support and advice.

## FUNDING

This work was supported by research grants from the Cystic Fibrosis Foundation; and National Institutes of Health [grant numbers COOPER21G0, GM127374] to B.S.C., National Institutes of Health; and National Science Foundation [grant numbers GM118139, CMMI–1548571] to Y.E.G. and P.C.N., and National Science Foundation [grant number PHY–2210452] to P.C.N.

## Conflict of interest statement

None declared.

## Supplementary Information

**Supplementary Figure 1:**
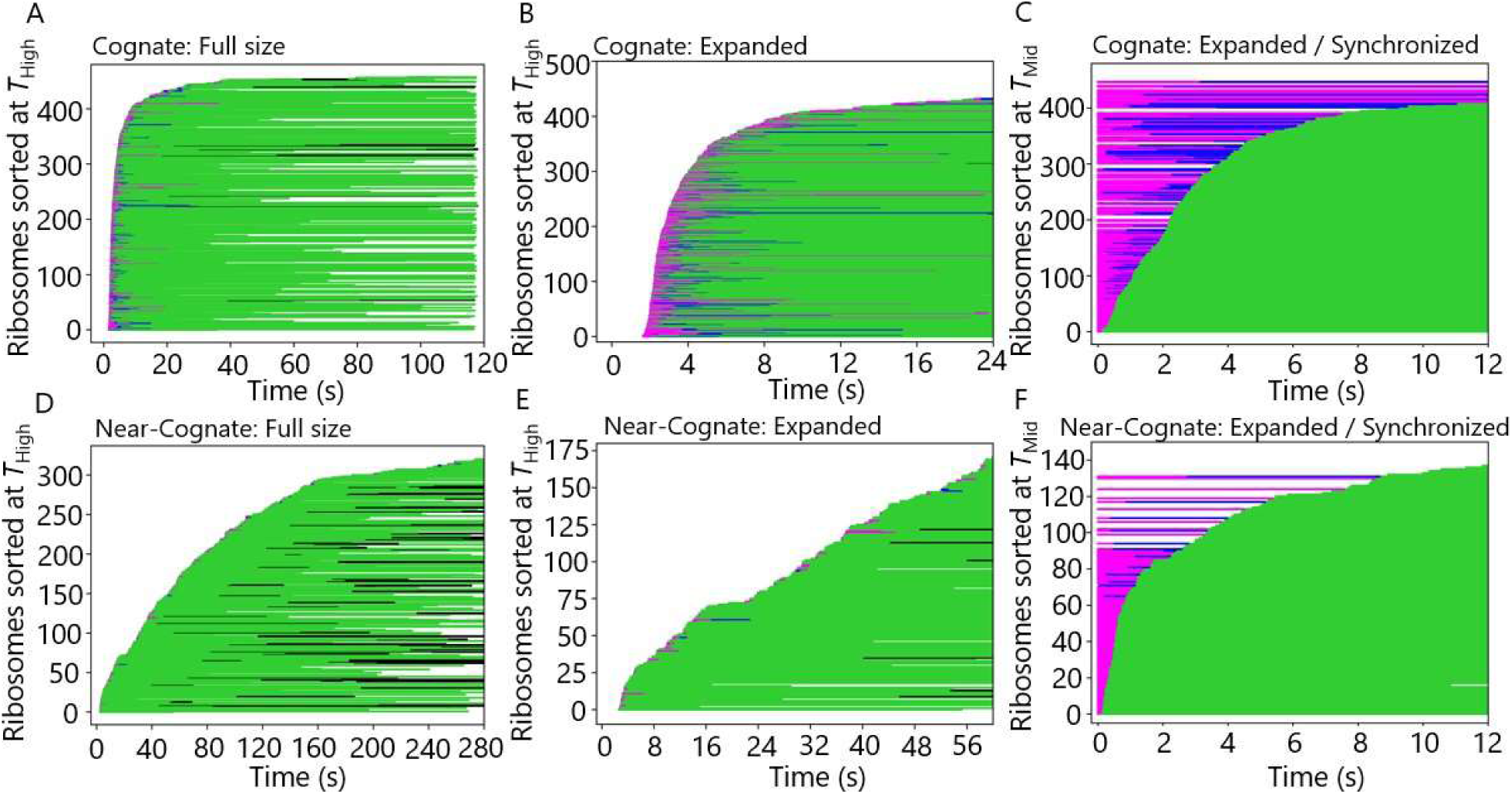
Rastergrams of series of traces for cognate and near-cognate ternary complexes. Each single-molecule FRET recording was categorized according to four different FRET states: Background FRET with or without sampling events prior to stable binding event (white), low-FRET transient state occurring upon stable binding (purple), high-FRET transient state upon stable binding (blue), mid-FRET state maintained for at least 10 s (green), and background FRET that occasionally followed stable binding but contains sampling events (black). The colored sequences were sorted according to the first arrival times of Cy5 FRET (TC binding), synchronized at the start of the movies and stacked vertically. A). Full-length single-molecule traces for cognate mRNA; B). As in (A) but with expanded time base; C). Color scheme as in (A) and (B) but sorted according to the time of reaching the mid-FRET state and synchronized (plotted t = 0) at the first arrival time of FRET. D), E) and F). As (A), (B), and (C), respectively except for near-cognate mRNA. All traces shown were obtained by injecting 25 nM Cy5-labeled ternary complex 1.0 second after the start of video recording.

**Supplementary Figure 2:**
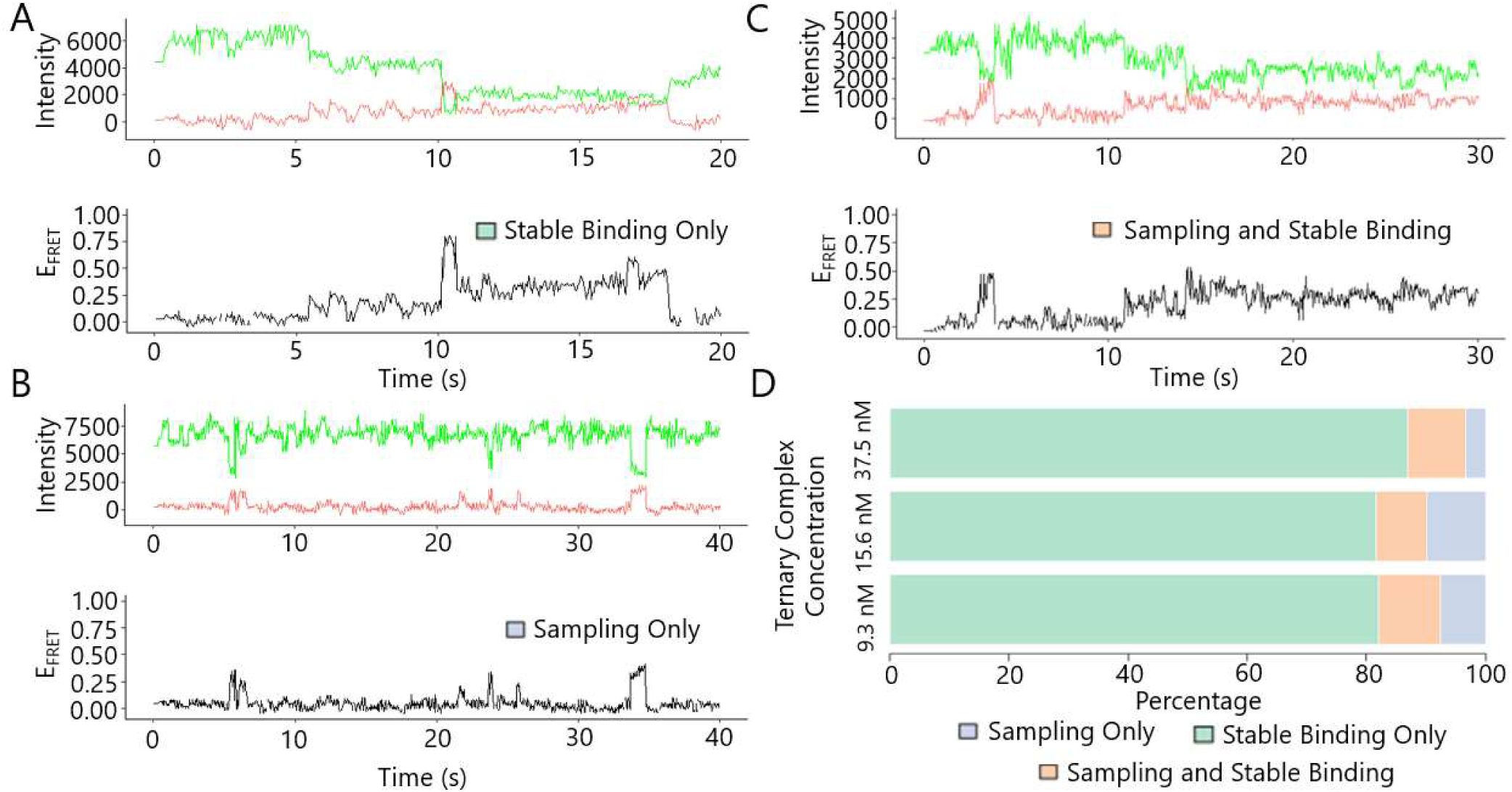
Classification of single molecule traces for cognate mRNA Measurements. A). Representative trace containing only stable binding. B). Representative trace containing only transient sampling events. C). Representative trace containing both a transient sampling event and stable binding. D). Stacked bar charts of the percentage of traces containing only stable binding events (green), sampling events and stable binding (orange), or only sampling events (blue). Data obtained at 9.3 nM, 15.6 nM and 37.5 nM TC combined.

**Supplementary Figure 3:**
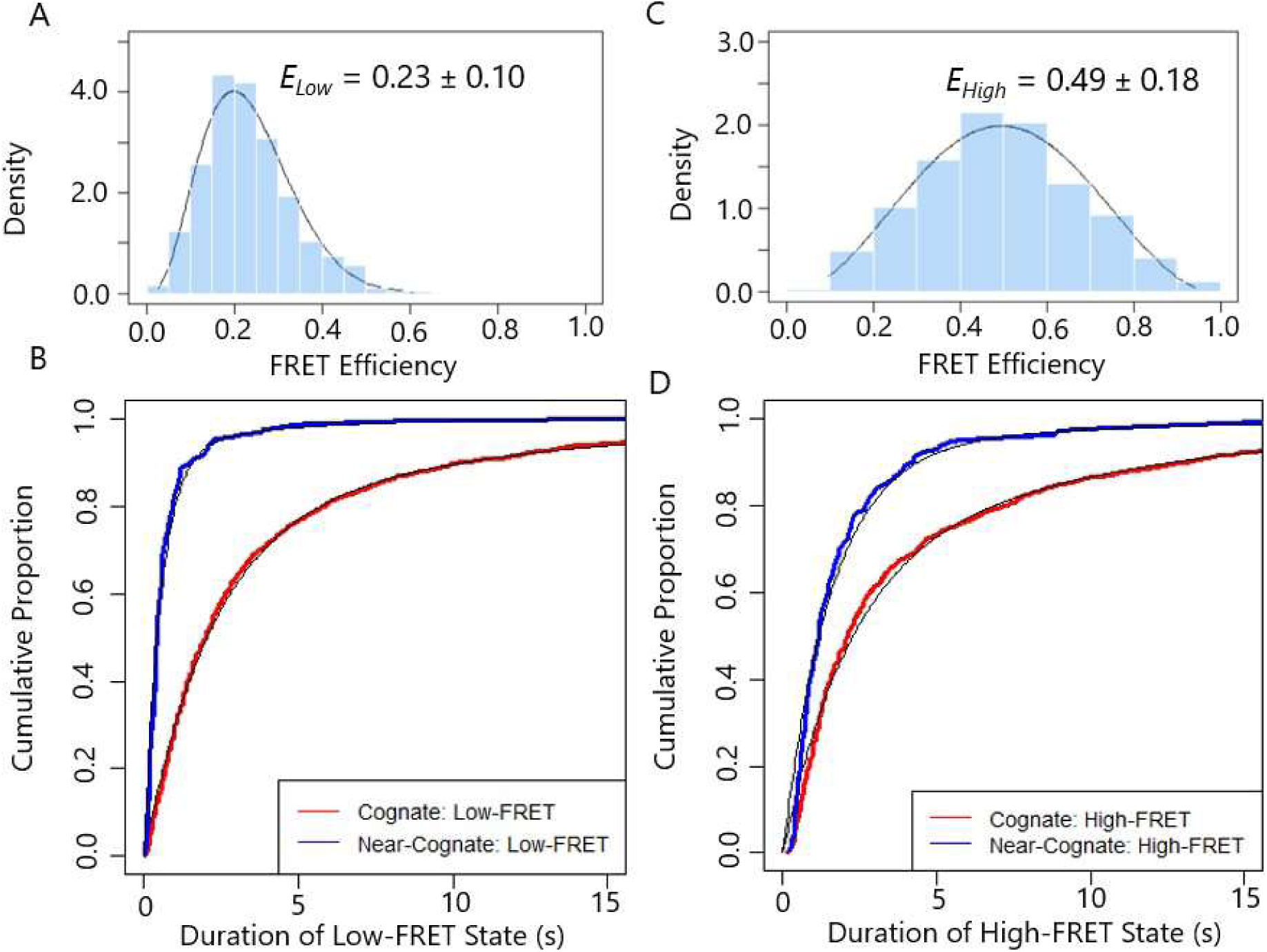
Lifetime and FRET efficiency analysis of ELow and EHigh states in stable binding of labeled TC to the ribosome. A). ELow FRET efficiency distribution for cognate and near-cognate mRNA. Black line is a fitted beta distribution. B). Cumulative distributions of lifetimes of ELow for cognate (red) and near-cognate (blue) mRNA. Black lines are fitted sums of two exponentials. C) and D). As in (A) and (B) respectively, but for the high-FRET state. Lifetimes and FRET efficiencies were generated from ≥ 200 traces per condition.

**Supplementary Figure 4:**
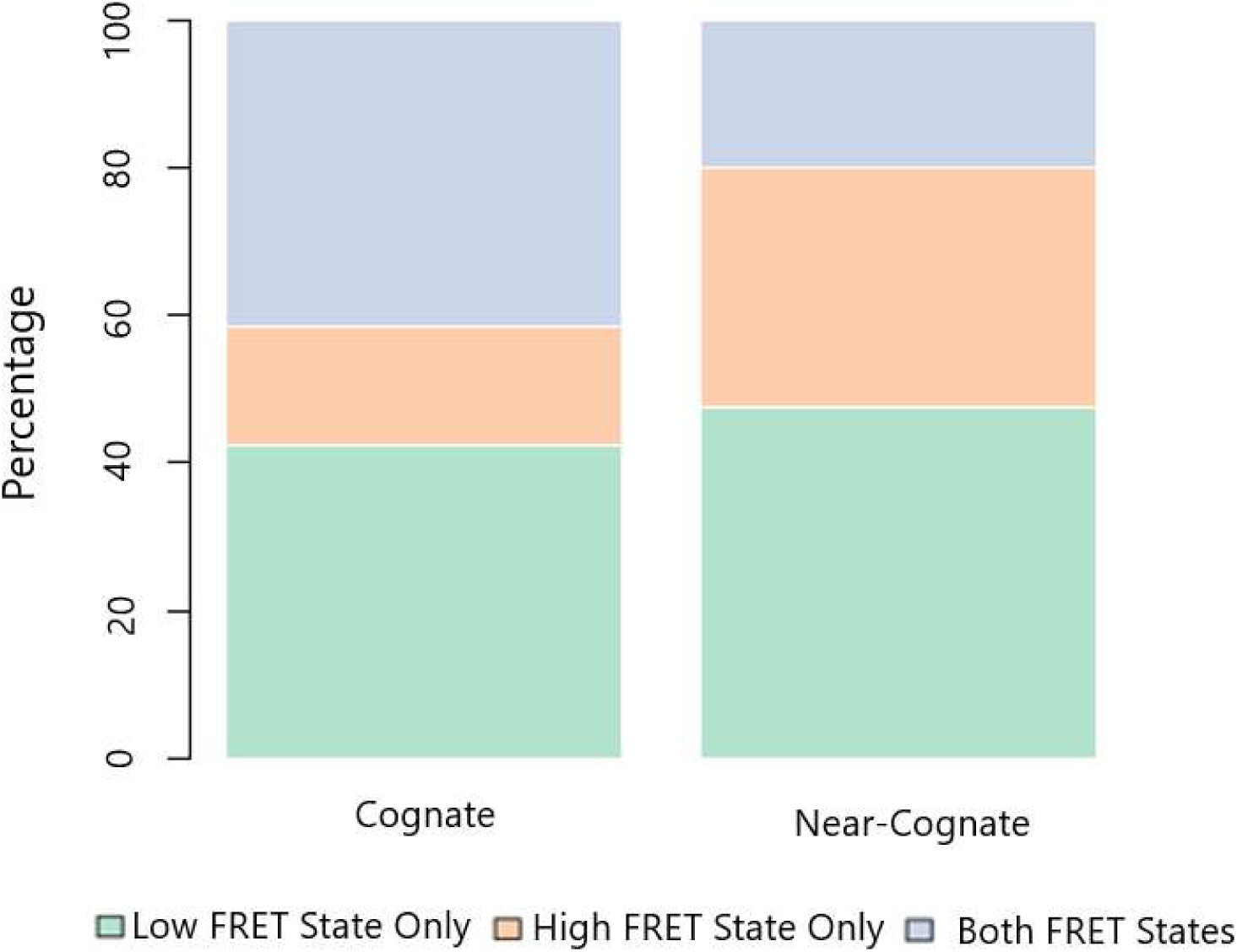
Classification of FRET states for cognate and near-cognate stable binding events containing the mid-FRET state. Preceding the mid-FRET state, recordings contained only the low-FRET state (green), only the high-FRET state (orange), or both (blue).

**Supplementary Figure 5:**
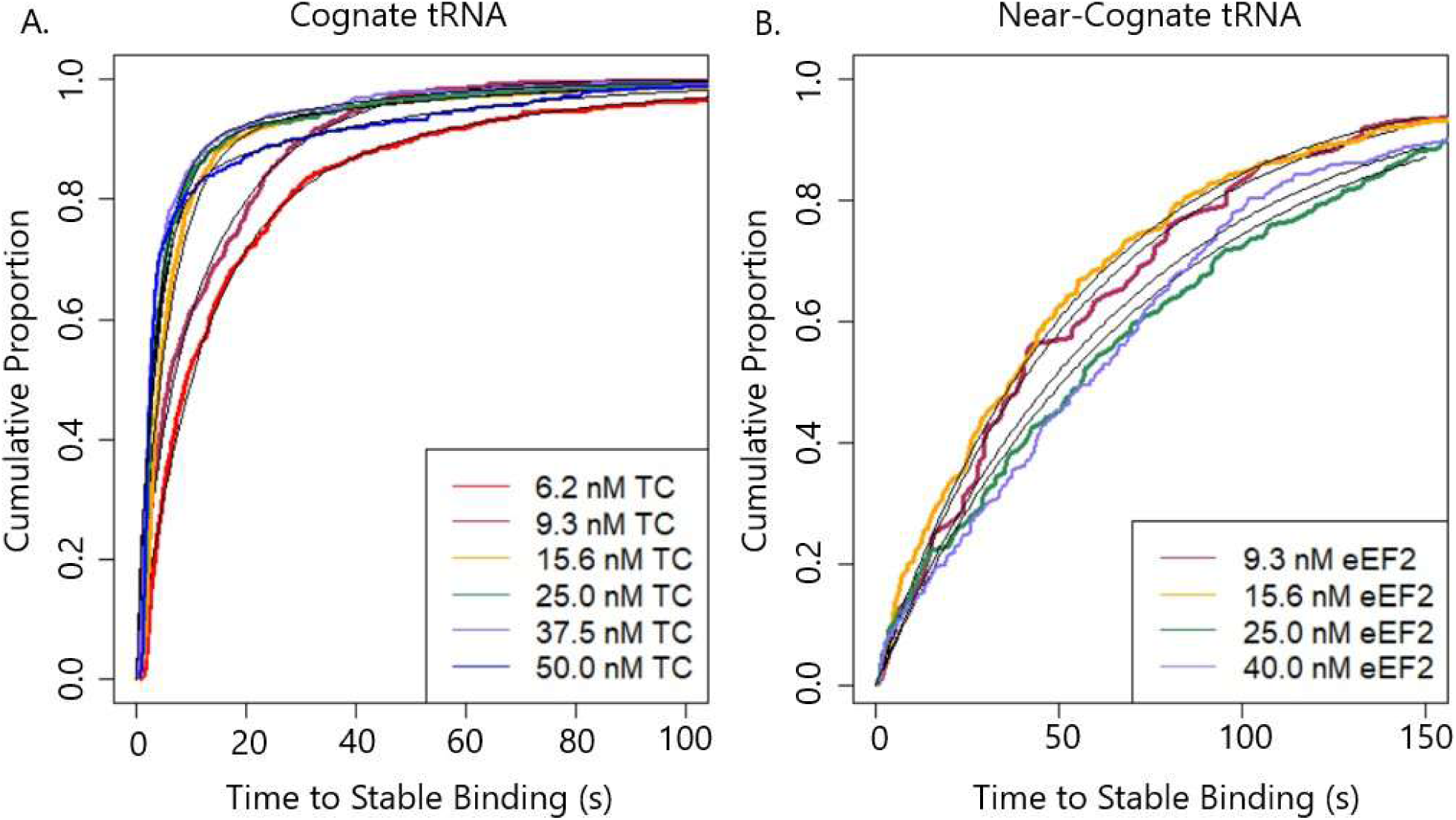
Time to stable binding for cognate and near-cognate mRNAs. A). Cumulative distributions of the time between delivery of Trp-TC (Cy5) and stable binding (TSB-All Traces) for the cognate condition. n ≥ 200 for each curve. Black lines are the sum of two exponential components fitted in MEMLET which also generated the rate constants plotted in Fig. 2C. B). Cumulative distributions of the time between delivery of Trp-TC (Cy5) and stable binding (TSB-All traces) for near-cognate binding. n ≥ 200 for each curve. Rate constants obtained by double-exponential fitting are plotted in Fig. 4D and E.

**Supplementary Figure 6:**
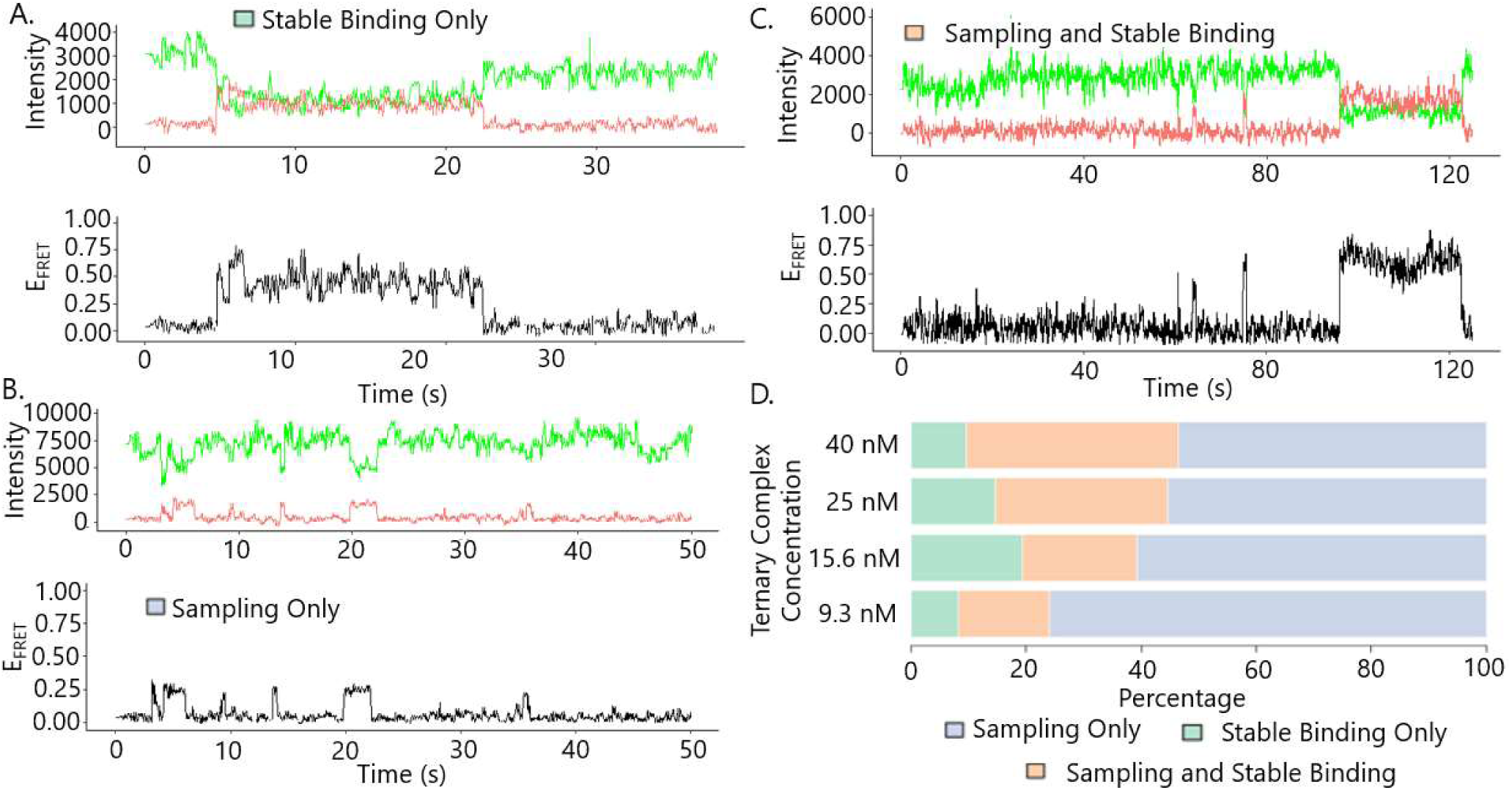
Classification of traces for near-cognate mRNA measurements. A). Representative single-molecule trace containing only a stable binding event; B). only transient sampling events; C). transient sampling and then stable binding. D). Stacked bar charts showing the percentage of traces of the three classes when 9.3, 15.6, 25, or 40 nM ternary complex was injected.

**Supplementary Figure 7:**
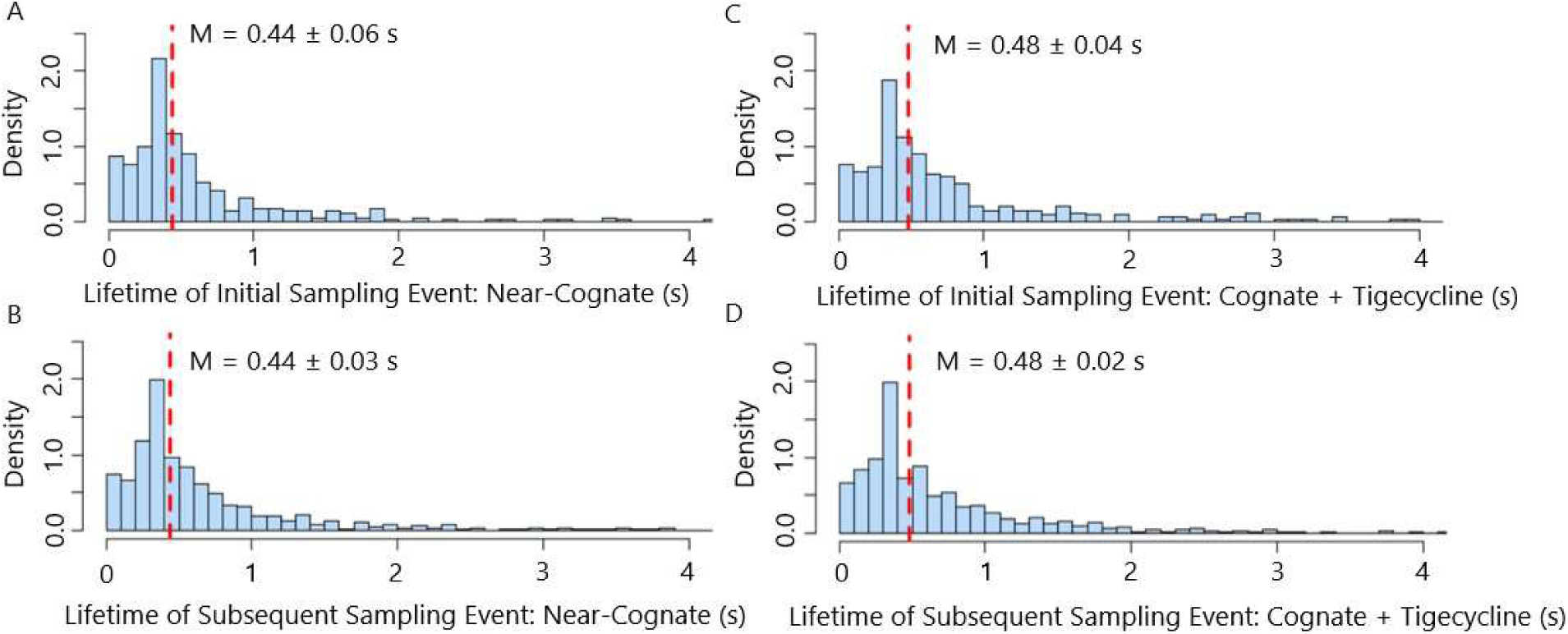
Histograms of lifetimes of sampling events A). Initial sampling event (τInitial) for near-cognate TC. B). Subsequent sampling events (τSubsequent) for near-cognate TC. C). Initial sampling event (τInitial) for cognate tRNA in the presence of 100 mM tigecycline. D). Subsequent sampling events (τSubsequent) for cognate tRNA in the presence of tigecycline. Red vertical dashed lines represent median values (M) for each histogram. Each histogram was generated from ≥400 traces and ≥1000 sampling events in total.

**Supplementary Figure 8:**
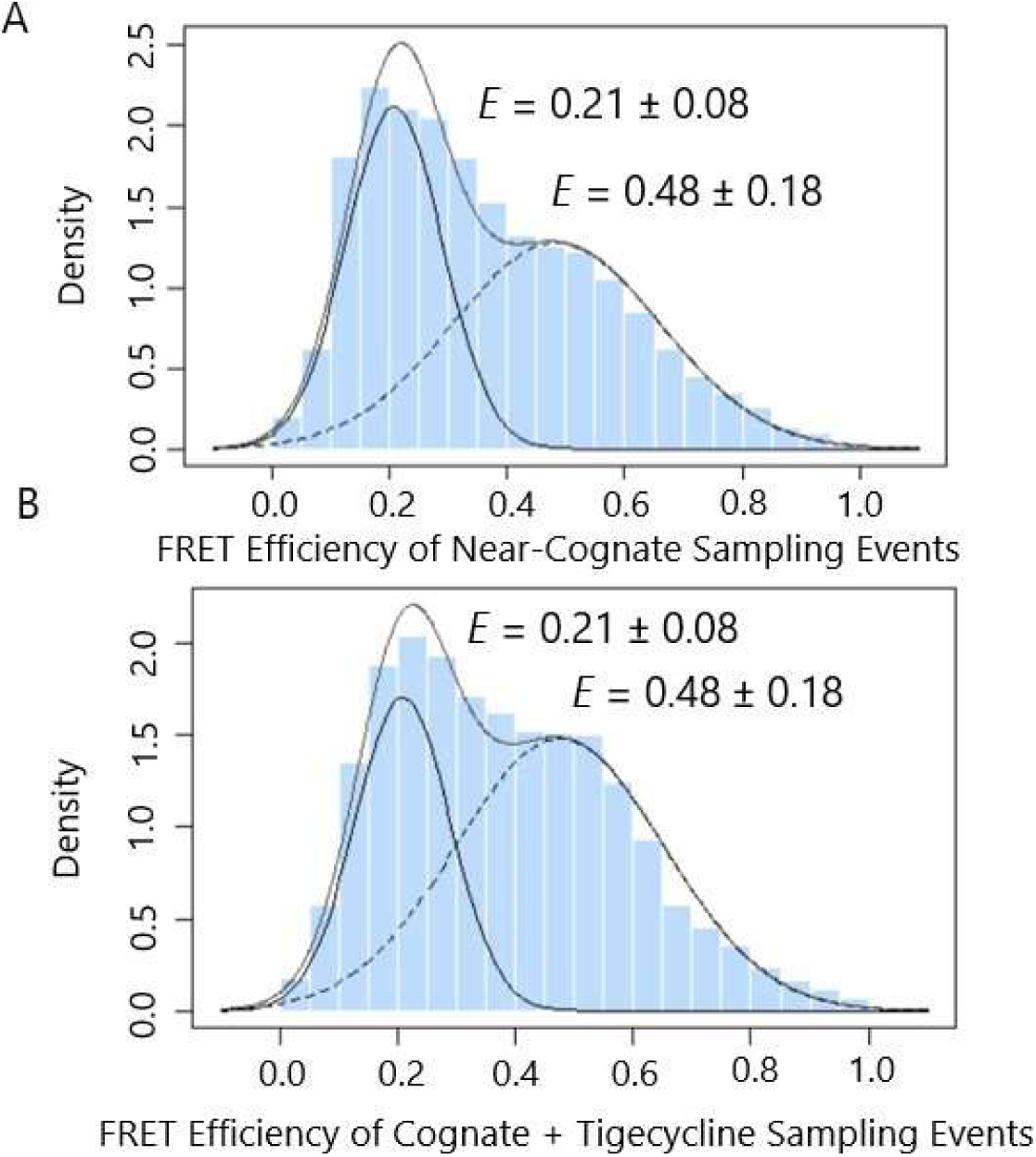
FRET efficiency distributions for Trp-TC (Cy5) sampling events of post-translocation ribosomes. A). near-cognate Trp-TC (Cy5). B). cognate Trp-TC (Cy5) in the presence of tigecycline. The distributions were fitted by the sums of two Gaussian components and were generated from ≥ 400 traces and ≥ 1000 sampling events in total.

**Supplementary Figure 9:**
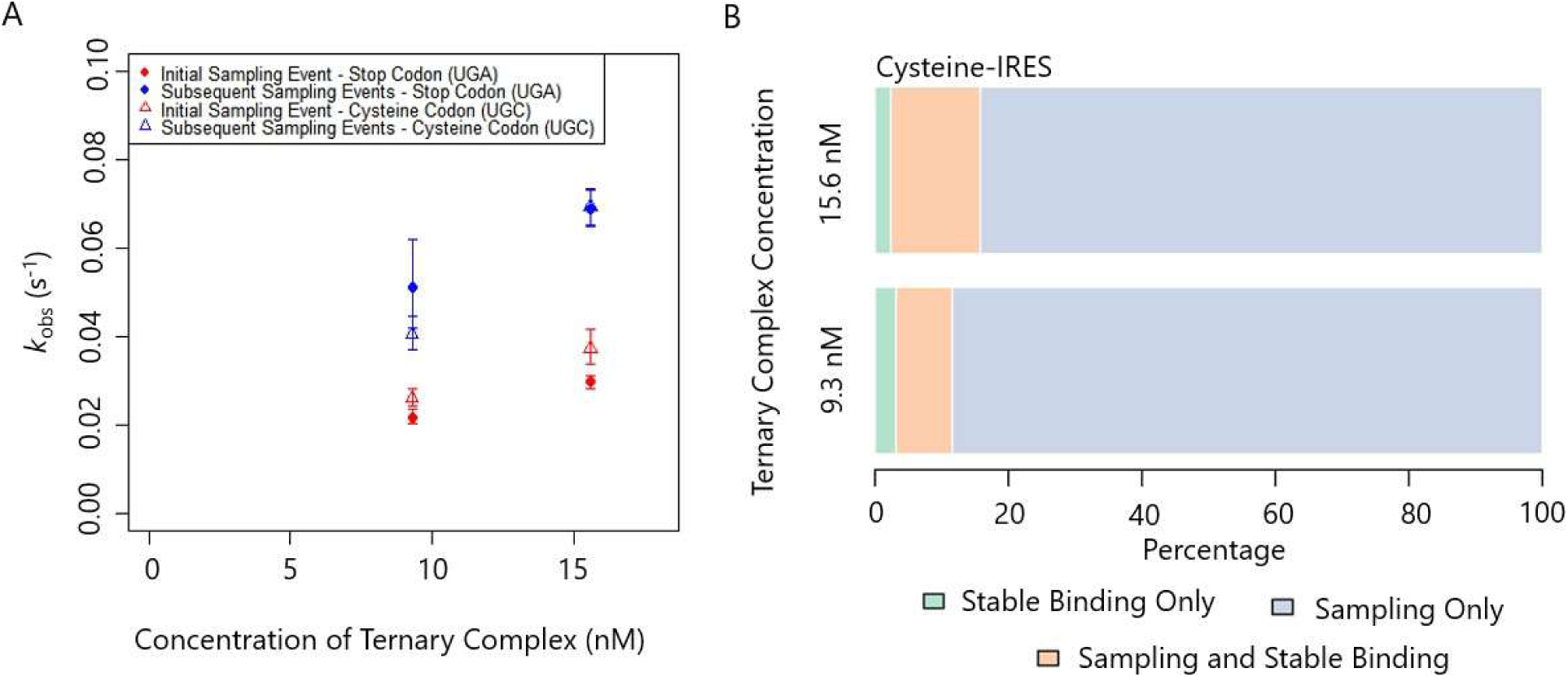
Binding kinetics for UGA (Cys) mRNA by Trp-TC (Cy5). A). Rate constants for initial (red symbols) and subsequent (blue) sampling events comparing UGA (stop, diamonds) and UGC (Cys, open triangles) codons. Error bars represent 95% confidence intervals generated in MEMLET by bootstrapping. Data for the stop mRNA are re-plotted from Fig. 3 D for comparison. B). Percentage of traces that contained only stable binding events (green), stable binding and sampling events(orange), and only sampling events (blue) at 9.3 and 15.6 nM ternary complex.

**Supplementary Figure 10:**
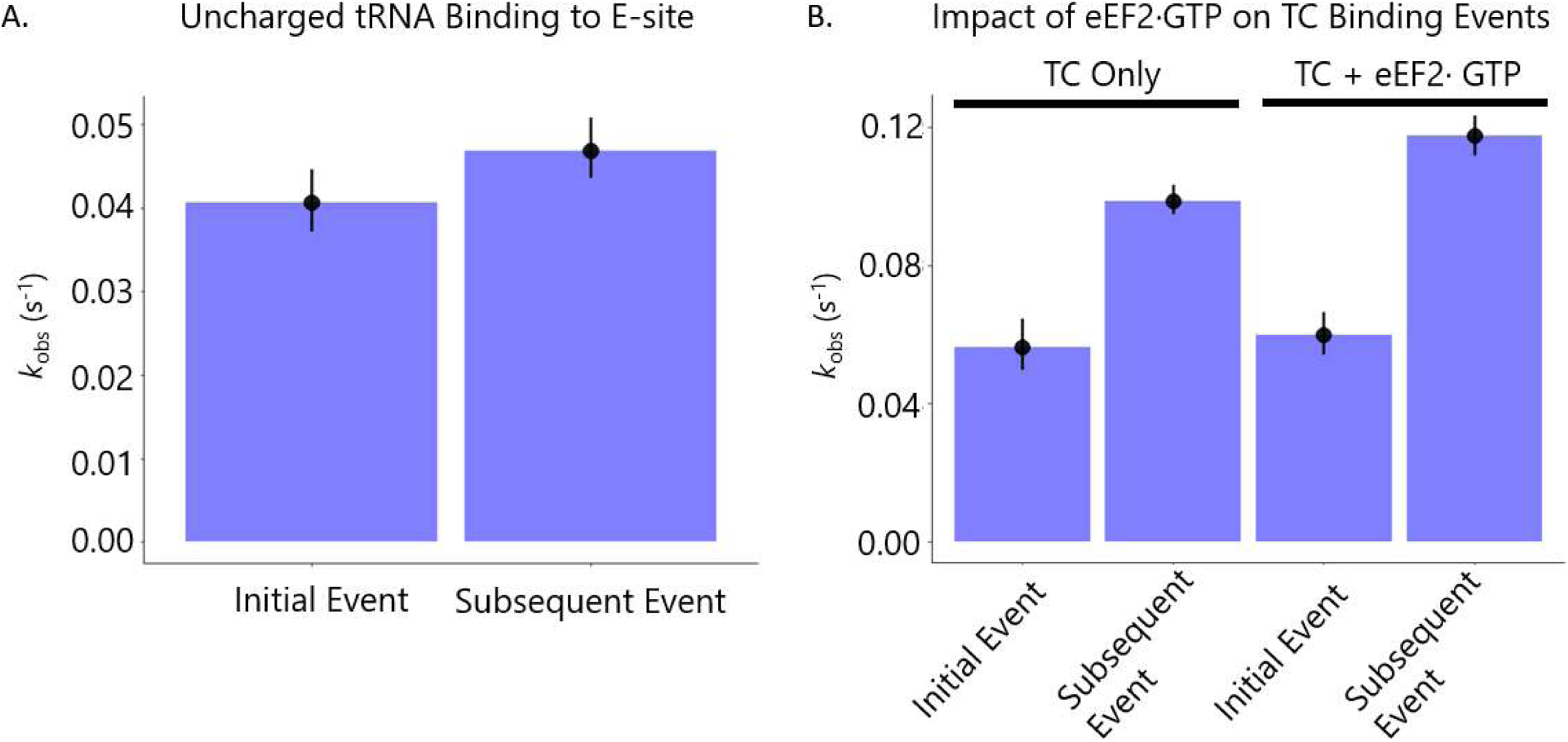
Tests for possible mechanisms for acceleration of near-cognate TC binding. A). Codon-independent binding of uncharged Cy5-labeled Trp-tRNA to the E-site of post-translocation ribosomes in the absence of eEF1A. Initial and subsequent sampling events at 9.3 nM are not significantly different. B). Effect of 5 mM eEF2·GTP on rate of initial and subsequent sampling events for near-cognate tRNA at 25 nM Trp-TC (Cy5). Rate constants for the initial and subsequent sampling events are similar in the presence or absence of eEF2·GTP presumably bound to the P1 stalk or other putative “channeling” binding sites. Error bars represent 95% confidence intervals obtained by bootstrapping.

**Supplementary Figure 11:**
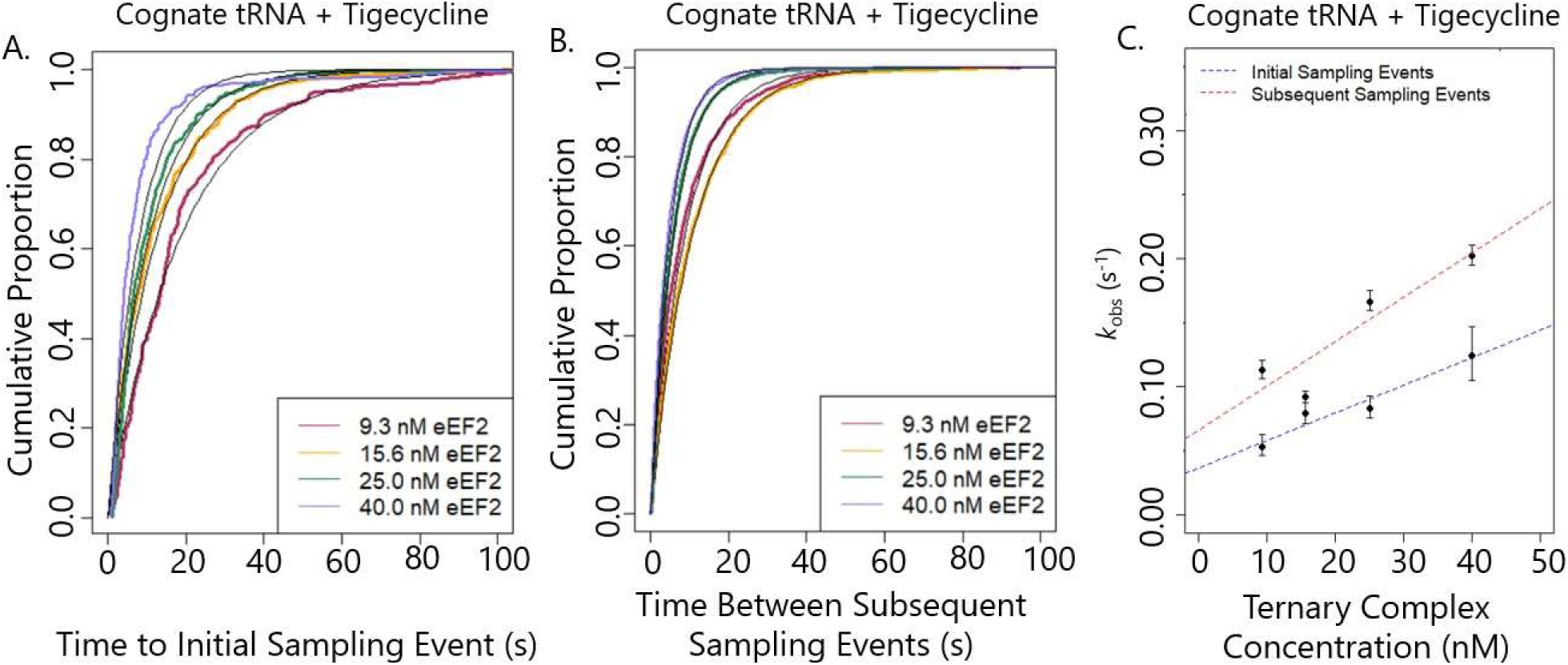
Dynamics of initial and subsequent sampling events for cognate mRNA in the presence of tigecycline. A). Cumulative distributions of the time between delivery of Trp-TC (Cy5) and initial sampling events (TInitial Binding) for cognate tRNA in the presence of tigecycline. Black lines represent single exponential fits. n ≥ 200 for each curve. B). As in A but for subsequent cognate sampling events (TSubsequent Binding) in the presence of tigecycline. n ≥ 500 for each curve. C). Rate constants for the initial and subsequent sampling events obtained by fitting single exponential functions to each lifetime distribution and plotted vs. TC concentration. Error bars represent 95% confidence intervals obtained by bootstrapping.

**Supplementary Figure 12:**
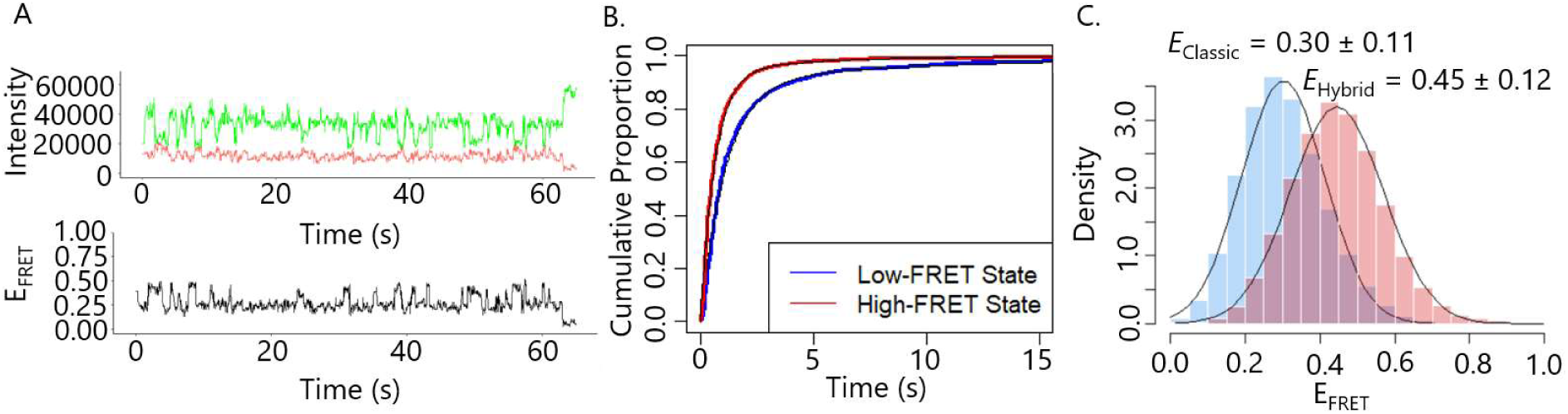
Dynamics of pre-translocation complexes. A). Representative single-molecule recording of low- and high-FRET fluctuations in ribosomes stalled in the PRE-translocation state. B). Cumulative distributions of the durations of the low- and high-FRET PRE states fitted by double exponential functions (black curves). C). Histograms of the FRET efficiency distributions of the low- and high-FRET states fitted by beta distributions. Cumulative distributions and histograms were generated from ≥500 traces with ≥2000 FRET transitions in total.

**Figure S13:**
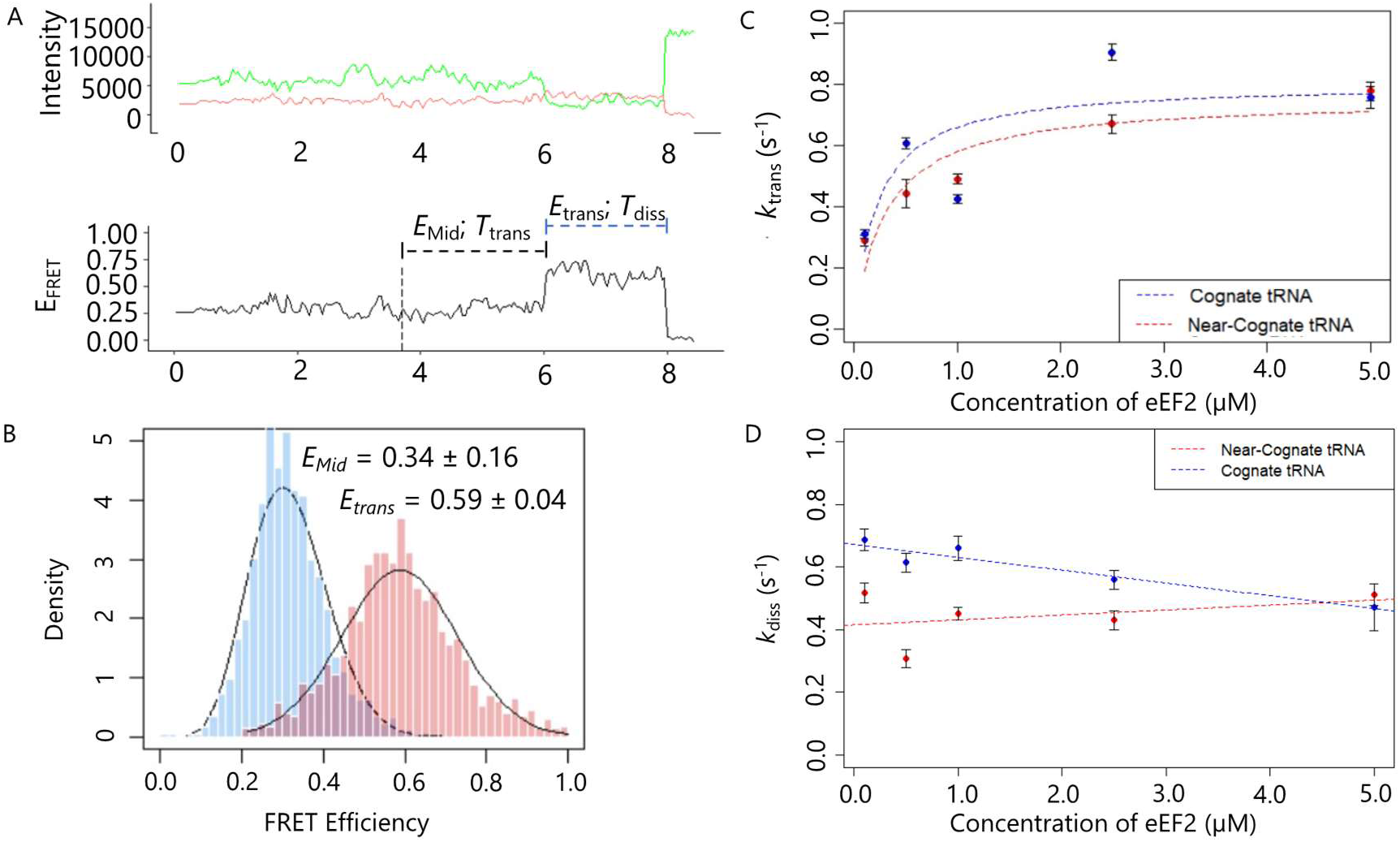
Kinetics of translocation. A). Representative single-molecule trace during translocation. Dashed line at 3.7 seconds shows the time at which eEF2 and GTP were delivered into the sample. Histograms of FRET efficiency distributions for the E_Mid_ portion of the trace and the E_trans_ state (shown in A) with fits of the beta distribution. C). Observed rate constants for the time from injection of eEF2·GTP to the formation of the E_trans_ state (T_trans_). Error bars represent standard error of the mean. D). Observed rate constants for the duration of the E_trans_ state following eEF2·GTP delivery (T_diss_). Error bars represent 95% confidence intervals. Measurements of translocation for the cognate tRNA condition were generated from ≥150 traces per condition. Measurements of translocation for the near-cognate tRNA using 0.1, 0.5, 1, 2.5 and 5 µM eEF2 were generated from 118, 51, 236, 97, and 134 traces, respectively.

**Supplementary Figure 14:**
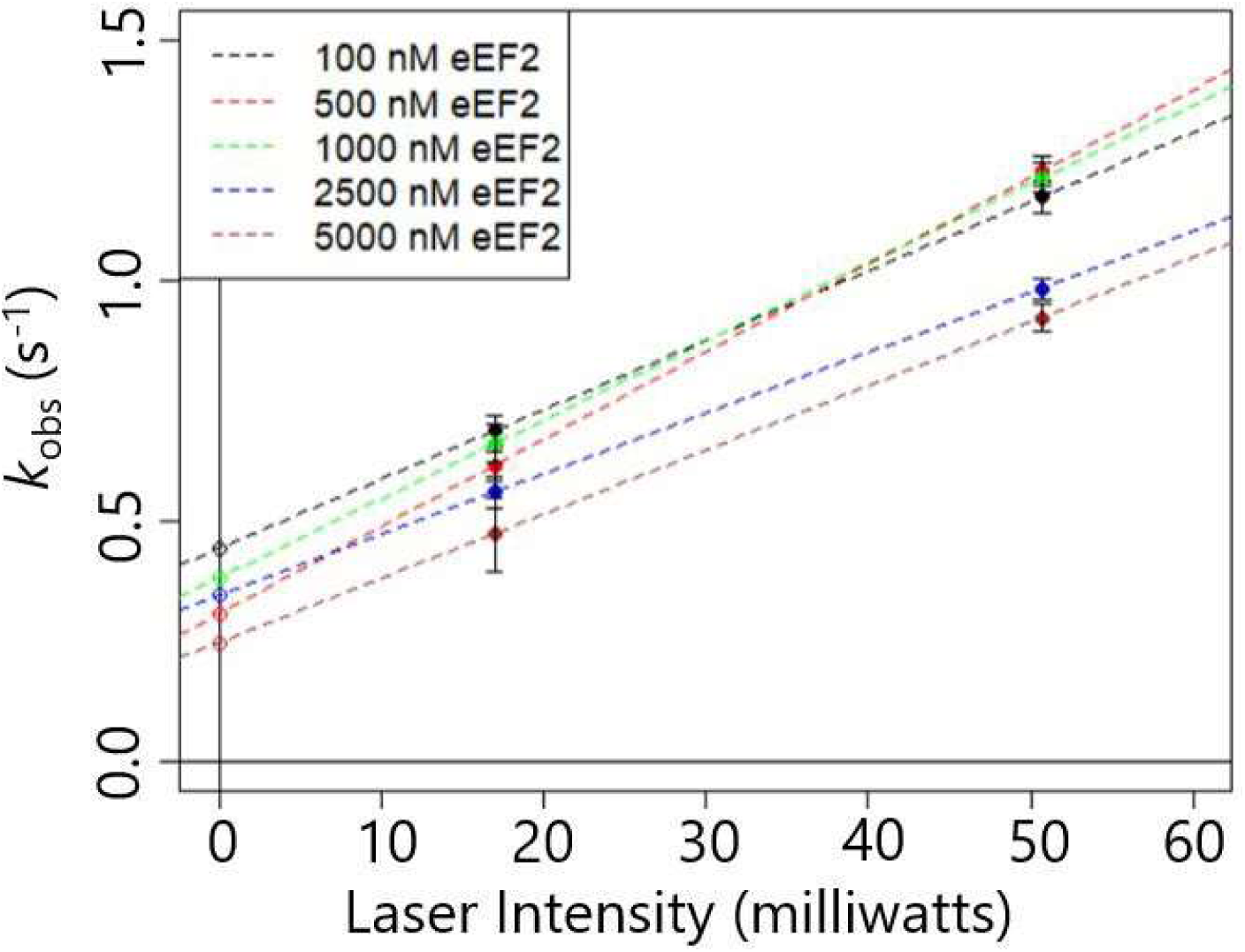
Rate constants for the duration of the high-FRET state following addition of eEF2·GTP for cognate tRNA at two different laser intensities to estimate the contribution of photobleaching. Photobleaching rate at laser intensity of 15.6 milliwatts (the power level for the main experiments) was estimated by extrapolating the lines determined by the measured rates. Error bars represent standard errors of the mean. Measurements were generated from ≥ 150 traces per condition.

**Table S1:** Parameters associated with stable binding of labeled tRNA to the ribosome for the cognate tRNA condition. TLow Binding is defined as the time from Trp-TC (Cy5) injection to ELow, THigh Binding is defined as the time from Trp-TC (Cy5) injection to EHigh and TMid Binding is defined as the time from Trp-TC (Cy5) injection to EMid. τLow is the lifetime of the ELow state while τHigh is the lifetime of the EHigh state. All parameters described above are illustrated in Fig. 2 A. Error bars represent the standard error of the mean.

| Table S1: Parameters associated with stable binding of labeled tRNA to the ribosome for cognate tRNA |  |  |  |  |  |
| --- | --- | --- | --- | --- | --- |
| Parameter | $T_{\text{Low Binding}} \text{ (s)}$ | $T_{\text{High Binding}} \text{ (s)}$ | $T_{\text{Mid Binding}} \text{ (s)}$ | $\tau_{\text{Low}} \text{ (s)}$ | $\tau_{\text{High}} \text{ (s)}$ |
| Condition (nM) |  |  |  |  |  |
| 6.2 | $19.2 \pm 1.7$ | $22 \pm 2$ | $31.2 \pm 1.5$ | $4.53 \pm 0.44$ | $6.19 \pm 0.57$ |
| 9.3 | $8.1 \pm 0.7$ | $10.1 \pm 0.8$ | $11.8 \pm 0.5$ | $4.45 \pm 0.31$ | $4.92 \pm 0.41$ |
| 15.6 | $6.8 \pm 1.0$ | $8.7 \pm 1.4$ | $11.01 \pm 0.74$ | $2.85 \pm 0.47$ | $3.32 \pm 0.69$ |
| 25 | $6.7 \pm 0.9$ | $11 \pm 1$ | $15.2 \pm 1.1$ | $4.62 \pm 0.52$ | $6.24 \pm 0.78$ |
| 50 | $5.3 \pm 1.2$ | $6.05 \pm 0.68$ | $13.32 \pm 0.97$ | $4.26 \pm 0.59$ | $4.52 \pm 0.49$ |

**Table S2:** Measurements derived from Cognate traces. A) Parameters associated with stable binding for traces that do not contain sampling prior to stable binding, traces that do contain sampling prior to stable binding or all traces that contain stable binding regardless of the presence of sampling events. TSB-No Sampling is defined as the time from injection of the Trp-TC (Cy5) until stable binding for traces that did not contain sampling events prior to stable binding, whereas TSB-With Sampling is defined as the time from injection of the Trp-TC (Cy5) until stable binding for traces that contained at least one sampling event prior to stable binding. TSB-All Traces is defined as the time from injection of Trp-TC (Cy5) until stable binding for all traces that contain stable binding regardless of the presence of prior sampling events. Tinitial Sampling is the time from injection to the initial sampling event in traces that contain at least one sampling event prior to stable binding and Tfinal Binding is the time from the last sampling event in a trace to the stable binding event. τInitial is the lifetime of the initial sampling event in traces that contain at least one sampling event prior to stable binding. Pa is defined as the proportion of traces that had stable binding events but no sampling events. Ns is the mean number of sampling events that occurred prior to stable binding for the different trace types. Error bars represent the standard error of the mean.

| Table S2: Measurements derived from Cognate Traces |  |  |  |  |  |  |  |  |  |
| --- | --- | --- | --- | --- | --- | --- | --- | --- | --- |
| Trace Identity | Stable Binding Only |  | Sampling and Stable Binding |  |  |  |  | All Stable Binding Traces |  |
| Parameter | $T_{\text{SB-No Sampling (s)}}$ | $P_a$ | $T_{\text{initial Sampling (s)}}$ | $\tau_{\text{initial (s)}}$ | $T_{\text{SB-With Sampling (s)}}$ | $T_{\text{Final Binding (s)}}$ | $N_s$ | $T_{\text{SB-All Traces (s)}}$ | $N_s$ |
| Condition (nM) |  |  |  |  |  |  |  |  |  |
| 9.3 | $12.3 \pm 0.5$ | 0.90 | $12 \pm 3$ | $0.57 \pm 0.04$ | $24 \pm 2$ | $12.0 \pm 1.2$ | $1.23 \pm 0.07$ | $12.3 \pm 0.6$ | $0.14 \pm 0.02$ |
| 15.6 | $9.9 \pm 0.2$ | 0.90 | $9 \pm 2$ | $0.57 \pm 0.04$ | $28 \pm 5$ | $9.8 \pm 1.9$ | $1.21 \pm 0.09$ | $9.9 \pm 0.6$ | $0.12 \pm 0.02$ |
| 37.5 | $7.3 \pm 0.2$ | 0.90 | $6 \pm 1$ | $0.57 \pm 0.04$ | $15 \pm 2$ | $6.0 \pm 1.1$ | $1.3 \pm 0.1$ | $7.3 \pm 0.6$ | $0.15 \pm 0.02$ |

**Table S3:** Parameters associated with Time to Intermediate Stable Binding States for Near-Cognate tRNA. TLow Binding is defined as the time from Trp-TC (Cy5) injection to ELow, THigh Binding is defined as the time from Trp-TC (Cy5) injection to EHigh and TMid Binding is defined as the time from Trp-TC (Cy5) injection to EMid. τLow is the lifetime of the ELow state while τHigh is the lifetime of the EHigh state. All parameters described above are illustrated in Fig. 2 A. Error bars represent the standard error of the mean.

| Table S3: Parameters associated with stable binding of labeled tRNA to the ribosome for near-cognate tRNA |  |  |  |  |  |
| --- | --- | --- | --- | --- | --- |
| Parameter | $T_{\text{Low Binding}} \text{ (s)}$ | $T_{\text{High Binding}} \text{ (s)}$ | $T_{\text{Mid Binding}} \text{ (s)}$ | $\tau_{\text{Low}} \text{ (s)}$ | $\tau_{\text{High}} \text{ (s)}$ |
| Condition (nM) |  |  |  |  |  |
| 9.3 | $48 \pm 9$ | $45 \pm 10$ | $60 \pm 6$ | $0.45 \pm 0.07$ | $1.6 \pm 0.3$ |
| 15.6 | $47 \pm 6$ | $43 \pm 5$ | $52 \pm 3$ | $0.62 \pm 0.07$ | $2.32 \pm 0.51$ |
| 25 | $83 \pm 8$ | $78 \pm 11$ | $74 \pm 4$ | $0.94 \pm 0.16$ | $1.47 \pm 0.20$ |
| 40 | $75 \pm 7$ | $69 \pm 7$ | $70 \pm 3$ | $1.18 \pm 0.46$ | $2.68 \pm 0.50$ |

**Table S4:** Measurements derived from Near-Cognate traces. A) Parameters associated with traces that contained only sampling events, stable binding events without prior sampling events, stable binding events with at least one prior sampling event and all traces that contain stable binding regardless of the presence of sampling events. TSB-No Sampling is defined as the time from injection of the Trp-TC (Cy5) until stable binding for traces that did not contain sampling events prior to stable binding, whereas TSB-With Sampling is defined as the time from injection of the Trp-TC (Cy5) until stable binding for traces that contained at least one sampling event prior to stable binding. TSB-All Traces is defined as the time from injection of Trp-TC (Cy5) until stable binding for all traces that contain stable binding regardless of the presence of prior sampling events. Tinitial Sampling is the time from injection to the initial sampling event and TSubsequent Sampling is the time between sampling events that occur after the initial sampling event. TInitial Sampling and TSubsequent Sampling are illustrated in Fig. 3 A. TFinal Binding is the time from the last sampling event in a trace to the stable binding event for traces that contain at least one sampling event prior to stable binding. τInitial is the mean lifetime of the initial sampling event and τSubsequent is the mean lifetime of subsequent sampling events. Pa is defined as the proportion of traces that had stable binding events but no sampling events. Ns is the mean number of sampling events that occurred prior to stable binding for the different trace types. Error bars represent the standard error of the mean.

| Table S4: Measurements derived from Near-Cognate Traces |  |  |  |  |  |  |  |  |  |  |  |
| --- | --- | --- | --- | --- | --- | --- | --- | --- | --- | --- | --- |
| Trace Identity | Sampling Only |  |  |  | Stable Binding Only |  | Sampling and Stable Binding |  |  | All Stable Binding Traces |  |
| Parameter | $T_{\text{Initial Sampling}}$ (s) | $T_{\text{Subsequent Sampling}}$ (s) | $\tau_{\text{Initial}}$ (s) | $\tau_{\text{Subsequent}}$ (s) | $T_{\text{SB-No Sampling}}$ (s) | $P_a$ | $T_{\text{SB-With Sampling}}$ (s) | $T_{\text{Final Binding}}$ (s) | $N_s$ | $T_{\text{SB-All Traces}}$ (s) | $N_s$ |
| Condition (nM) |  |  |  |  |  |  |  |  |  |  |  |
| 9.3 | 46 ± 2 | 20 ± 2 | 0.53 ± 0.02 | 0.47 ± 0.01 | 40 ± 5 | 0.63 | 74 ± 11 | 19 ± 5 | 1.53 ± 0.15 | 57 ± 6 | 0.58 ± 0.10 |
| 15.6 | 34 ± 1 | 14.5 ± 0.4 | 0.53 ± 0.02 | 0.47 ± 0.01 | 30 ± 3 | 0.53 | 68 ± 5 | 18 ± 2 | 2.18 ± 0.11 | 54 ± 3 | 1.02 ± 0.08 |
| 25 | 24 ± 1 | 7.9 ± 0.2 | 0.53 ± 0.02 | 0.47 ± 0.01 | 30 ± 5 | 0.25 | 79 ± 5 | 15 ± 1 | 6.8 ± 0.5 | 73 ± 4 | 5.08 ± 0.52 |
| 40 | 20 ± 1 | 5.5 ± 0.1 | 0.53 ± 0.02 | 0.47 ± 0.01 | 26 ± 7 | 0.18 | 7 ± 4 | 11 ± 1 | 8.2 ± 0.6 | 68 ± 4 | 6.70 ± 0.56 |

**Table S5:** Comparison of rate constants determined in this study versus rate constants determined in previous study by Ng et al (20). All experiments conducted in the current study were done at 24 °C, whereas all experiments conducted in Ng et al. were done at 37 °C.

| Table S5: Comparison of rate constants determined in this study versus rate constants determined in Ng et al. |  |  |  |  |
| --- | --- | --- | --- | --- |
| Rate Constant | Data Source | Cognate | Near-Cognate | Cognate / Near-Cognate |
| $k_{\text{Pep}}$ | Ng et al. | $0.3 \text{ s}^{-1}$ | $0.06 \text{ s}^{-1}$ | 5 |
| $k_{\text{Mid Binding}}$ | Current Study | $0.1 \text{ s}^{-1}$ | $0.02 \text{ s}^{-1}$ | 5 |
| $k_{-1}$ | Ng et al. | $2.2 \text{ s}^{-1}$ | $2.3 \text{ s}^{-1}$ | 1 |
| $k_{\text{Initial}}$ | Current Study | $2.0 \text{ s}^{-1}$ | $2.0 \text{ s}^{-1}$ | 1 |
| $k_1$ | Ng et al. | $52 \mu\text{M}^{-1}\text{s}^{-1}$ | $10 \mu\text{M}^{-1}\text{s}^{-1}$ | 5.2 |
| $k_{\text{SB}} - \text{Cognate tRNA}$<br>$k_{\text{Subsequent Sampling}} - \text{Near-Cognate tRNA}$ | Current Study | $15 \mu\text{M}^{-1}\text{s}^{-1}$ | $4.4 \mu\text{M}^{-1}\text{s}^{-1}$ | 3.4 |

Global maximum-likelihood analysis of near-cognate binding

Single-molecule fluorescence resonance energy transfer yields rich datasets that can be used to interrogate the mechanisms of the eukaryotic ribosome. Rather than reduce such data to first moments, for example, mean waiting times as functions of tRNA concentration, here we sought a principled, maximum-likelihood approach that objectively fits the entire distributions of measured waiting times for a proposed kinetic model. Our approach is computationally inexpensive and amenable to bootstrap estimation of errors. Briefly, we found expressions for the likelihood functions for certain FRET transitions, then maximized likelihood by using a customized approach similar to [1].

Our experiment discriminates between states with zero FRET signal and three different nonzero levels. The resulting datasets give the times of transitions between those FRET levels. Referring to the proposed kinetic scheme (Figure 7), note that:

- States **1**, **2**, **F**, and **E** are experimentally indistinguishable “zero FRET” states.
- The transition from **3** to **4** is rapid, and sometimes the dwell in **3** is missed, so in this discussion we will lump these states together as a combined “low or high” FRET state.
- **R** also has low FRET.
- State **6** has a distinct FRET level indicating final binding.

Rather than fit to the very general state scheme (Figure 7), we made further simplifications. In addition to combining states **3** and **4** as just mentioned, we:

- Neglected the possible direct transition **5** → **F**.
- Neglected the possible direct transition **3** → **E**.

All binding reactions will be assumed to follow simple mass-action. Thus, we introduce the abbreviation *κ*_1_ for the pseudo-first order rate constant *k*_1_[*TC*], where [*TC*] is concentration of ternary complex, and similarly *κ*_8_ = *k*_8_[*TC*], *κ*_6_ = *k*_6_[*TC*]. In addition, define the branching fractions *β* = *k*_4_*/*(*k_−_*_2_*_a_* +*k*_4_), and *α* = *k*_7_*/*(*k*_7_ + *κ*_6_).

### 1 Data

First we identified state transitions by the means described in Materials and Methods. This procedure yielded many series of transition times:

- *t*_1_, time from injection to initial sampling event binding (if any). This is a transition zero FRET→low or high fret that is followed later by a transition back to zero FRET. *t*_1_ is an abbreviation for the quantity called *T*_InitialSampling_ in the main text.
- *d*_1_, duration of initial sampling event until it ends with unbinding.

Subsequent binding events (transitions from zero to low or high FRET after the first unbinding) were further subdivided into two classes:

- If the rebinding was itself sampling (i.e., followed later by unbinding), then the waiting time is an instance of *t*_2_, an abbreviation for the quantity called *T*_SubsequentSampling_ in the main text.
- If the rebinding was final (i.e., followed later by a transition to state **4**), then the waiting time is an instance of *t*_3_, an abbreviation for the quantity called *T*_FinalSampling_ in the main text.

Finally, every ribosome that eventually ended in stable binding gave an instance of a discrete random variable:

- *N*_samp_, the number of unbinding events prior to stable binding, or 0 if stable binding occurs on the first attempt.

### 2 Predicted distributions

The family of models under consideration allowed us to write down the predicted probability distributions of observed data analytically, as described below. We then maximized likelihood over the unknown parameter values holding the experimental data (and known parameters like [*TC*]) fixed. Exhaustive grid searches for the maxima were used to avoid the possibility of finding local (not global) maxima; however, the likelihood surfaces were inspected, and each was found retrospectively to be well behaved.

#### 2.1 First binding

This event is defined as the first transition out of zero-FRET after injection of TC.

**Probability density function** In the model, initial binding is a Michaelis– Menten process:

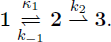

(As mentioned, we assume an irreversible second step, or *k_−_*_2_ = 0.) Let

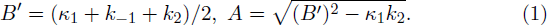

Note *B^′^ >* 0 and 0 *< A < B^′^*. The probability density function (PDF) for completion times is [2]

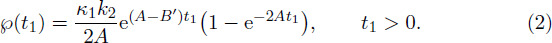

This function is always positive, as it must be. It can be used to compute a likelihood function that determines *k*_1_, *k_−_*_1_, and *k*_2_ from *t*_1_ data at several values of [*TC*]. Fig. S14 upper left compares our global fit to our data.

For the initial *sampling* binding events, we must randomly reject a constant fraction *β* of these events (because that fraction proceed to stable binding), but that complication has no effect on the PDF of times.

**Mean rate** Eqn. (2) implies that the mean rate is

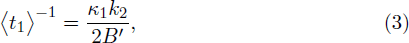

which agrees with the classical Michaelis–Menten formula [2]. This function of [*TC*] is shown as the red curve in Fig. S14 lower left.

**Cumulative density function** The CDF is

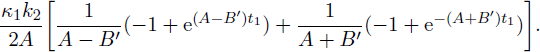

#### 2.2 First unbinding

This event is defined as the first transition back to zero FRET (if any) after initial binding. The binding duration *d*_1_ should be distributed as a single exponential, because the first unbinding is definitely from **3**. The PDF in our model is therefore 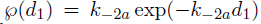, independent of [*TC*]. The CDF is 1 − e*^−k−^*^2^*^ad^*^1^ .

We measured separate sets of *d*_1_ values for each of various [*TC*]. To maximize log likelihood, average over them all:

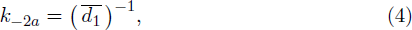

where overbar denotes sample mean.

Fig. S14 top right shows that the distribution of *d*_1_ values starts exponential, but has a long tail. We interpreted this by concluding that the transition to **4** isn’t literally permanent; there can be unbinding from **5** after several seconds, which should not be considered unbinding for our purposes.

Accordingly, we dropped the few events with *d*_2_ *>* 4 s in the maximum-likelihood fit.

#### 2.3 Subsequent sampling

“Subsequent binding” is defined as the first transition from zero-FRET to low or high-FRET after first unbinding. It can therefore occur either by a return to state **3** via an undetected sojourn in **E**, or by transition to the dead-end state **R**. “Subsequent sampling” is a more restricted class of those subsequent binding events that are followed by unbinding (that is, it excludes those transitions to **3**/**4** that proceed to stable binding). We now work out the probability distribution for first passage to either **3** or **R** given that the event is sampling.

Let us enumerate every possible outcome. Whenever the ribosome arrives at **R**, it must eventually return to **F**. Call these events of type *α*. But from **3**/**4** there is probability *β* of passing on to **5** (type *β* event) and 1 − *β* of returning to **F** (type *γ* event). Types *α, β, γ* are mutually exclusive and one of them must eventually happen in our model if we start from **F**.

Integrating the three PDFs *℘_α_*, *℘_β_*, and *℘_γ_*, then adding them, does yield 1.

Then the PDF for **F** to transition to low or high-FRET at time *t*, then return to **F**, is:

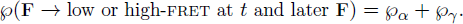

The conditional probability density is this function of *t* divided by P(later **F**). But it’s easier just to observe that it must be a normalized PDF, and to compute the needed normalization constant.

Let

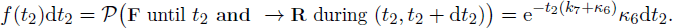

Next, consider

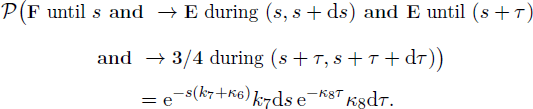

Let

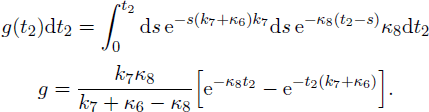

The PDF of *all* subsequent binding times is *f* + *g*. The PDF of all *sampling* rebinding times is instead

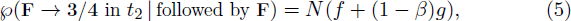

where the normalization factor is:

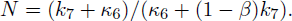

Fig. S14 center left compares our global fit to our data.

The CDF is *N* (*F* + (1 − *β*)*G*), where

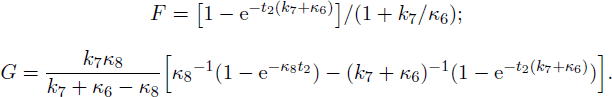

The mean waiting time for subsequent sampling is then

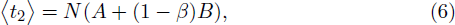

where

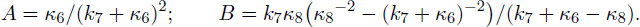

The mean rate is the reciprocal of the mean waiting time. Eqn. (6) defines a function of [*TC*], whose reciprocal is plotted as the blue curve in Fig. S14 lower left.

#### 2.4 Stable rebindings

For those binding events that proceeded to **5**, we need

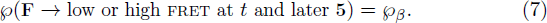

The conditional probability density is this function of *t* divided by P(later **5**). Again it’s easier just to observe that it must be a normalized PDF, and to compute the needed normalization constant.

Thus, the PDF of all stable rebinding times is

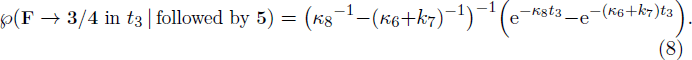

Note that *β* drops out of this expression; once we specify that the next transition is to **5**, the branching fraction for the last step is immaterial.

The PDFs Eqns. (5) and (8) can be used to compute a joint likelihood function that determines *k*_8_, *k*_6_, *k*_7_, and *β* from *t*_2_ and *t*_3_ data.

The expectation of waiting time for stable rebinding from Eqn. (8) is

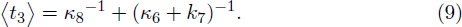

Eqn. (9) defines a function of [*TC*], whose reciprocal is plotted as the green curve in Fig. S14 lower left.

### 2.5 Number of sampling attempts

We next wish to find the model’s prediction for the discrete probability distribution P_samp_ of the number of sampling attempts before stable binding (*N*_samp_). Again define *α* and *β* as above. After initial binding, there is a branch, with probability to proceed directly to stable binding (zero sampling events):

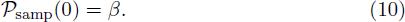

Eqn. (10) defines a (constant) function of [*TC*] that is shown in Fig. S14 lower right.

If the system instead falls through to a zero-FRET state, then it arrives in **F**. Each time it arrives there, it can:

- Enter **R** (probability 1 − *α*), which guarantees another visit to **F** prior to stable binding, or:
- Proceed to **E** (probability *α*) and thence to **3**/**4**, where we face a further branch:

**–** “Succeed” (pass to **5**), or

**–** “Fail” (return to **F**).

Thus,

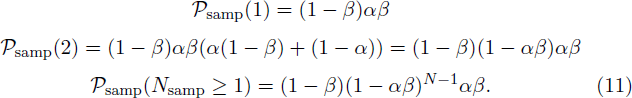

The family of discrete distributions just found can be used to compute a likelihood function that determines *β* from *N*_samp_ data, holding *α* fixed to the value found from fitting samplign rebindings.

Fig. S15 below shows our model’s predictions of P_samp_ as curves corresponding to Eqns. (10–11), and compares to our data. In a nutshell, P_samp_ *is a geometric distribution beyond N*_samp_ = 0, making it easy to obtain *α* and *β* via maximum likelihood. We can also compute the expectation

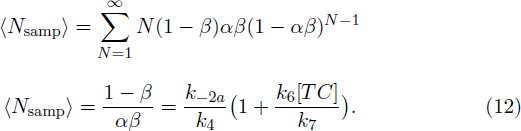

Eqn. (12) defines a function of [*TC*] that is shown as the curve in Fig. S14 lower center.

### 3 Best fits

Figs. S14 and S15 show that a single global fit successfully reproduces many different features of our experimental data. The best-fitting parameter values from this procedure are given in the table below.

Note that:

- Every individual data point contributed to its likelihood function. Thus, reduced statistics (such as those in the three graphs in the bottom row) were not given any privileged status.
- The lower-left curves are Eqns. (3) and the (reciprocal of) (6) with parameters taken from the maximum-likelihood fit.
- The *N*_samp_ curve (lower center) is Eqn. (12) with parameters taken from the maximum-likelihood fit.

**Figure S15:**
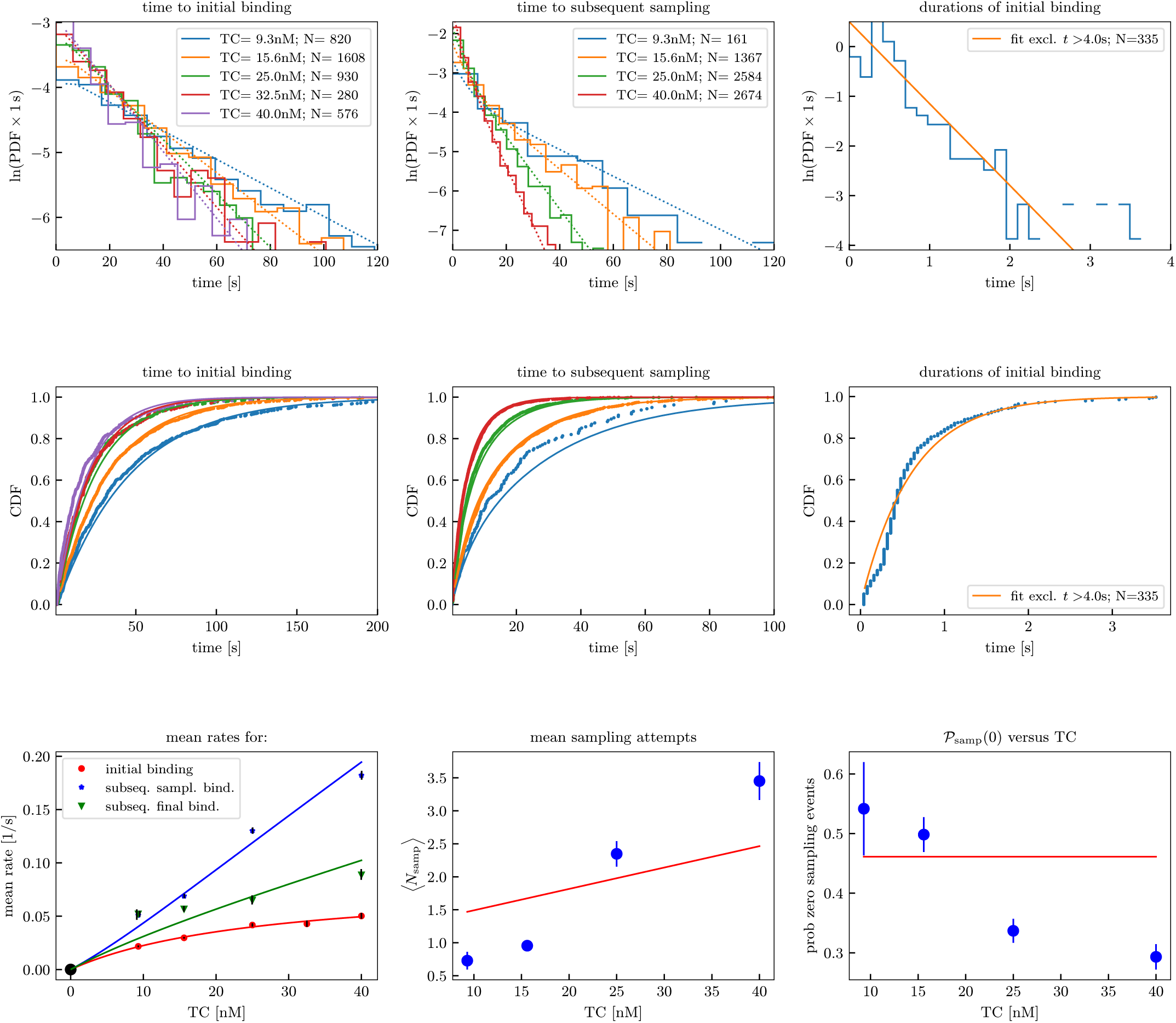
Global fit to all initial binding, subsequent sampling, and stable rebinding time distributions, and to the distribution of sampling attempt number. Smooth curves are analytic predictions (Sect. 2), based on the global fit values given in Sect. 3. Staircase plots are estimates of PDFs based on binned experimental data. Ragged curves on CDF plots are estimates from experimental data. Dots on the lower row were obtained by taking indicated averages of our experimental data. Error bars are derived from standard error of mean for experimental data.

**Figure S16:**
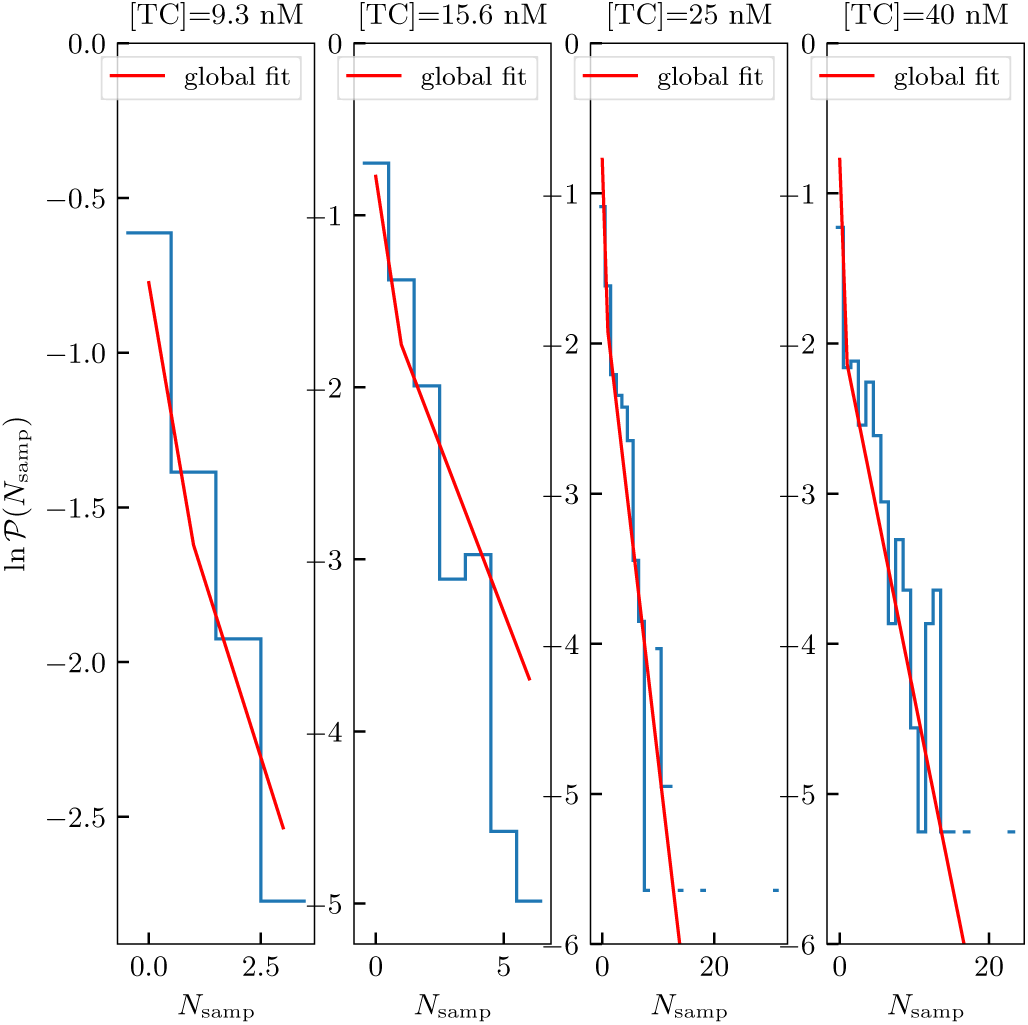
P(*N*_samp_) estimated from data compared to global best fit. Lines are predictions based on maximum-likelihood fit values given in Sect. 3, using Eqns. (10) and (11). Dots are the distributions estimated from experimental data.

In Fig. S15, lines again show the analytic predictions made earlier with best-fitting parameters. Symbols are estimated probabilities from experimental data. The experimental distributions do look like the analytic prediction, for example, the semilog plots are roughly straight lines for *N*_samp_ *>* 0.

**Fit values** Fitting was performed by iterating the preceding likelihood maximization steps until convergence. After best-fitting parameter values were formed, bootstrap replica datasets were created by sampling the data with replacement, and for each such choice a new set of maximally likely parameters was chosen. We found covariance matrices for the resulting clouds of points in parameter space, and used their eigenvalues to estimate standard deviations for the parameters. The individual values of *k*_1_ and *k_−_*_1_ are poorly constrained by the data, but their ratio (equilibrium dissociation constant) is well defined, as indicated in the table. The values shown are in reasonable agreement with those in the main text, which were obtained by fitting the mean rates and attempt numbers in a stochastic simulation of the model to the corresponding experimental values.

|  |  |
| --- | --- |
| $k_1$ | $21 \pm 2 \mu\text{M}^{-1} \text{s}^{-1}$ |
| $k_{-1}$ | $0.49 \pm 0.05 \text{s}^{-1}$ |
| $k_{-1}/k_1$ | $23 \pm 1 \text{nM}$ |
| $k_2$ | $0.083 \pm 0.008 \text{s}^{-1}$ |
| $k_{-2a}$ | $1.65 \pm 0.09 \text{s}^{-1}$ |
| $k_8$ | $3.6 \pm 0.1 \mu\text{M}^{-1} \text{s}^{-1}$ |
| $k_6$ | $4.5 \pm 0.1 \mu\text{M}^{-1} \text{s}^{-1}$ |
| $k_7$ | $0.16 \pm 0.01 \text{s}^{-1}$ |
| $k_{-2a}/k_4$ | $1.17 \pm 0.05$ |

## Notes

### Competing Interest Statement

The authors have declared no competing interest.

